# Semi-supervised segmentation of RNA 3D structures using density-based clustering

**DOI:** 10.1101/2025.01.12.632579

**Authors:** Quoc Khang Le, Eric Angel, Fariza Tahi, Guillaume Postic

## Abstract

A growing body of evidence shows that the biological activity of RNA molecules is not only due to their primary and secondary structures, but also to their spatial conformation. This is analogous to proteins, where investigating function, folding, or evolution often requires dividing the three-dimensional (3D) structure into subparts that can be studied individually. These independent substructures, known as protein “3D domains”, are geometrically defined as compact and spatially separate regions of the polypeptide chain. In RNA macromolecules, however, and to the best of our knowledge, no equivalent 3D-based concept has yet been formulated.

We present RNA3DClust, an application of the Mean Shift clustering algorithm to the RNA 3D structure partitioning problem. For this work, a dedicated post-clustering procedure was developed to address the peculiarities of delimiting 3D domains in RNA conformations. Tuning and benchmarking RNA3DClust required us to create reference datasets of RNA 3D domain annotations and to devise a new scoring function—the Chain Segment Distance (CSD)—for assessing segmentation quality. Importantly, we show that the domain decompositions produced by RNA3DClust are consistent with those based on RNA biological function and evolution. Finally, the emerging interest in long non-coding RNAs (lncRNAs) and their likeliness of containing folded regions has motivated us to generate an additional reference dataset of lncRNA predicted conformations. The resulting delineations of 3D domains by RNA3DClust illustrate the potential of our method for analyzing lncRNA 3D structures. Source code and datasets are freely available for download on the EvryRNA platform at: https://evryrna.ibisc.univ-evry.fr.

**GRAPHICAL ABSTRACT:** 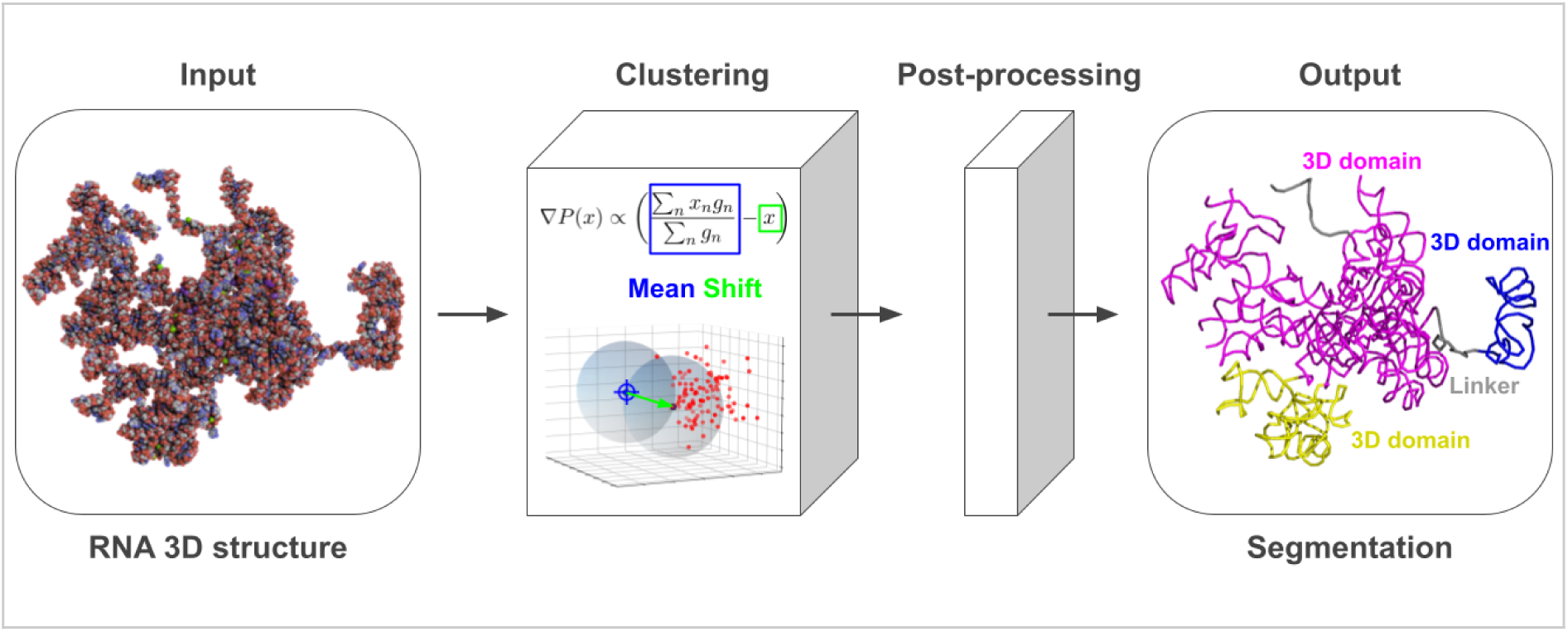

## 1. INTRODUCTION

Analysis is defined as the separation of a whole into its constituent parts in order to gain a better understanding of it. In the context of structural biology, it consists of dividing macromolecules into simpler yet informative substructures, which can be studied individually. In the case of proteins, the spatial conformation of a polypeptide chain, which is integral to its biological function, can be described through different levels of analysis that include, hierarchically: (i) secondary structures, which are local folding patterns that emerge from interactions between nearby amino acids along the protein backbone, (ii) supersecondary structures, which are made of several adjacent secondary structure elements, and (iii) 3D domains—sometimes referred to as “structural domains”—which are regions of the proteins that can fold, function or evolve independently. These latter subparts contribute to the dynamic nature of the protein, with “domain motion” being the process of 3D domains moving relative to each other, while keeping their individual structures. Domains, rather than entire protein structures, are also responsible for mediating interactions between multi-domain proteins [1]. All these biological properties make the question of how to parse domains critical for understanding the molecular mechanisms that operate in the cell, and for potential applications, such as protein engineering or drug design. Importantly, protein 3D domains are the basic units used in structure classification systems, such as the CATH [2], SCOP [3], and ECOD [4] databases.

In practice, 3D domains are central to the divide-and-conquer strategies employed for both *in vitro* determination and *in silico* prediction of protein structures. Experimental biologists often divide protein chains into autonomous folding units that are easier to purify and crystallize than the full-length macromolecule and yield higher-resolution crystallographic data. A single polypeptide chain can adopt multiple biologically active conformations due to inter-domain flexibility and motion. Therefore, rather than directly predicting the whole conformation, computational biology approaches predict the domain folds individually, before positioning them to compose the global structure. Successful methods such as AlphaFold [5] and RoseTTAFold [6] implicitly account for information on 3D domain organization, through their attention mechanism and architectural layers, respectively. Recently, it has been shown that the success rate of AlphaFold-Multimer [7] on multimeric structures can be raised by ∼20%, when adding a proper delimitation of the protein 3D domains to the input multi-sequence alignments [8]. Finally, it must be mentioned that, since the 15th edition of the CASP competition, 3D domains are partitioned to define evaluation units (EUs), which are used for both performance assessment and difficulty classification [9].

The structural organization of RNA macromolecules also exhibits multiple levels: next to the RNA sequence itself, the lowest is the secondary structure primarily formed by complementary base pairing—yet it is enriched and stabilized by a diverse array of non-canonical interactions. Higher in the hierarchy, the RNA motifs [10–12] are small, recurring secondary structure elements that act as modular building blocks of the overall RNA 3D shape. Examples include kink-turns, A-minor motifs, and tetraloop-tetraloop receptors. Beyond this, some RNAs can fold into intricate 3D structures driven by long-range interactions between motifs. Like the protein structure-function relationship, this tertiary level of structure dictates some RNAs’ biological functions [13], such as the catalytic activity of ribozymes, the molecular recognition of ribosomes, or the binding of specific ligands by aptamers to trigger regulatory responses. Besides, studying RNA tertiary structure is important from an evolutionary standpoint, as it supports the RNA world hypothesis by demonstrating RNA’s capacity to serve as both genetic information storage and catalyst, in early life forms [13].

Despite the fundamental significance of RNA 3D structure, the amount of experimental data available remains very limited in the reference databases, namely the Protein Data Bank (PDB) [14] and the Nucleic Acid Knowledgebase (NAKB) [15]. Consequently, the concept of protein 3D domain has not yet been transposed to RNA molecules (see Section 2.1). However, the investigation of RNA 3D domains is essential to implement the aforementioned divide-and-conquer strategies aimed at experimentally determining or computationally predicting the conformation of RNA chains—especially when they are long. Furthermore, this missing level of structural analysis in RNA will only become more detrimental over time, with the recent emergence of long non-coding RNAs (lncRNAs). This type of RNA molecule was initially seen as non-functional, until researchers unveiled their critical roles in regulating various cellular processes and their association with various diseases, including cancer, neurological disorders, cardiovascular diseases [16]. The minimum length of lncRNAs is defined as 200 nucleotides, but these long transcripts typically reach thousands of nucleotides, which make them likely to contain folded regions, akin to protein 3D domains.

Here, we propose the first computational study of RNA 3D domains, which are defined as compact and separate regions of the macromolecule—analogously to the original definition introduced by D. Wetlaufer for globular protein domains [17]. Thus, we considered different clustering methods taking as input the atomic coordinates of RNA 3D structures, and compared their capacity to find relevant domain limits (*i.e.* cluster boundaries). We chose this approach rather than adapting one of the existing protein-specific methods, because of the fundamental differences between peptide and nucleic acid chains, particularly with respect to flexibility. Furthermore, to account not only for the spatial distribution of atoms but also for the sequence continuity of the entire RNA molecule, we developed a dedicated post-clustering procedure. This work required us to create datasets of 3D domain annotations for the hyperparameter tuning and benchmarking procedures as, to the best of our knowledge, none has ever been published. As the ground truth, we used the visual compactness and separation criteria, as well as data about biological function taken from the literature. In this way, we have developed a semi-supervised clustering-based approach named RNA3DClust, which adapts the well-established Mean Shift algorithm [18] to the specificities of RNA 3D domains. Source code and datasets are freely available for download at: https://github.com/LeKhang97/RNA3DClust.

## 2. METHODS

### 2.1. Definitions

In the present study, we delineate compact and spatially separate regions of the RNA 3D structure. Since we adapted this seminal definition of 3D domains from proteins [17] to RNAs, several adjustments were necessary and are presented here. This section also describes the atomic representations used, as well as the distinction between 3D domains and other types of RNA substructures produced by previously published methods. An illustration of the RNA 3D domain concept is shown in Fig. 1.

**Figure 1.**
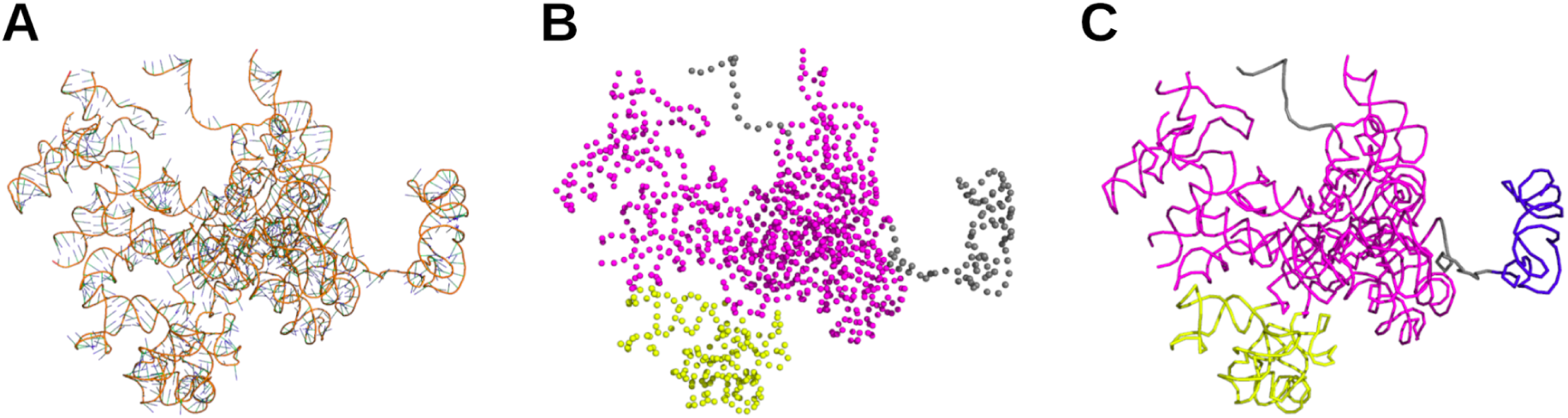
Definition of RNA 3D domains, illustrated with the LSUb ribosomal RNA (rRNA) from *Trypanosoma brucei brucei* (PDB entry: 8OVA, chain BB). (**A**) Cartoon representation of the RNA molecule. (**B**) Delimitation of 3D domains (shown in magenta and yellow), based on their compactness and spatial separation, and a sequence length ≥ 30 nt; each nucleotide is represented by its C3’ atom (spheres). (**C**) Helical regions, although less compact, are also annotated as 3D domains (blue). In this ribbon representation, the remaining unfolded segments shown in gray correspond to flexible “linker” regions.

#### 2.1.1. RNA 3D domains

For this initial definition of RNA 3D domains, we restrict the concept to single nucleic acid chains, as domains in proteins are always defined within individual polypeptide chains. Consequently, we do not consider regions that appear compact and separate only when multiple RNA chains are taken together. We set a minimum chain length of 30 nt. This threshold was based on the hammerhead ribozyme (HHR), which is the shortest and most studied RNA with a catalytic activity. Although shorter synthetic variants have been engineered, the natural form of the HHR is 43 nt. This provides an estimate of the minimum chain length required for a stable and functional RNA 3D structure. Additionally, we considered the fact that PIWI-interacting RNAs (piRNAs) exert their protein-binding function in a sequence-specific manner (*i.e.* independent of any particular RNA 3D fold), while being rarely longer than 32 nt. The same applies to small interfering RNAs (siRNAs) and microRNAs (miRNAs), which are typically ∼21–24 nt in length. Based on these observations, we established an arbitrary and rounded threshold of 30 nt for RNA 3D domains. This choice aligns with a similar convention in proteins, for which it is generally accepted that no stable fold is possible below ∼30 amino acids.

In addition to domains, 3D structures may also contain unfolded segments known as “**linkers**” in proteins, as they connect adjacent 3D domains. They provide flexibility, enable movement, and ensure proper spacing between domains, thereby playing crucial roles in domain-domain communication and conformational changes during function. Here, for RNA linker regions, we set a minimum sequence length of 10 nt. This threshold value effectively excludes the possibility of RNA structuration, as a stem-loop requires at least 3 unpaired nucleotides in the loop (to allow the backbone to bend) and a stem of 2–3 base pairs to fold and remain stable. When two 3D domains are connected by less than 10 nt, each half of the segment is assigned to the respective adjacent 3D domain; such RNA regions are not classified as linkers, as they are too short to confer meaningful flexibility. Unlike 3D domains, we also imposed a maximum sequence length for linkers, since longer RNA segments are less likely to remain unfolded. We empirically set this upper limit at 100 nt (see Section 3.3). Of note, we also call linkers unfolded RNA segments located at the 5’ or 3’ ends of the sequence, even though they do not *link* any pair of domains.

Finally, the base pairing between complementary RNA segments often results in the formation of double helices. Although not as compact as 3D domains, RNA **double helical regions** are still structured, which excludes them from the definition of linkers provided above. Therefore, to maintain consistency with the binary classification used in protein 3D analysis (*i.e.*, the “domain versus linker” distinction), we chose to include double helical regions within the definition of RNA 3D domains. It should be noted that helicity can also arise in single-stranded RNA, from base stacking alone (*i.e.*, without base pairing), particularly under high-salt conditions or in crowded environments. However, given their unstable and transient nature, we consider these single helical conformations part of the unstructured category, rather than 3D domains.

#### 2.1.2. Single-atom representations

The geometrical criteria of compactness and spatial separation, which originally defined 3D domains in proteins, rely on the (*x*, *y*, *z*) Cartesian coordinates of the macromolecule’s atoms. Here, to segment RNA 3D structures in an automated manner, we based our approach on clustering these atomic positions. As the resulting domain boundaries must be identified by their positions in the RNA sequence, we used a single-atom representation, *i.e.* selecting a single atom of each nucleotide. This approach parallels protein domain parsing programs that use either the Cα [19] or Cβ atoms [20] of the protein structures. For this work, we tested representations using either RNA’s C4’, C3’, or C1’ atoms, as they are part of the sugar-phosphate backbone and, therefore, present in all four residue types. Notably, the C4’ atom is used in multiple studies to approximate the center of the nucleotide without using base atoms (see [21] for a review). The C3’ atom is also the recommended and default option in structural alignment algorithms like RNA-align [22] and US-align [23]. The C1′-only model is less common in the literature: the positioning of the C1’ atom, connected directly to the base, can indeed lead to non-uniform spacing along the RNA backbone and make distance-based methods unreliable. Nevertheless, we included this third single-atom model in our study.

#### 2.1.3. Disambiguation

This article introduces the concept of parsing spatially separate and compact regions within RNA 3D structures. These 3D domains should not be confused with substructures from prior studies, which defined RNA decompositions based on different criteria and for different purposes. For example, the structural fragments used in tools such as RNA FRABASE 2.0 [24], Rosetta FARFAR [25], or FARFAR2 [26] are typically shorter than those produced by our method, which imposes a minimum sequence length of 30 nucleotides (see Section 2.1.1). With its size, spatial separation, and compactness, a 3D domain is expected to correspond to an RNA region capable of folding independently. In contrast, shorter fragments like FARFAR’s are designed to be assembled into the larger 3D structure of such an autonomous folding unit. As a result, an RNA structure may be composed of dozens or hundreds of such short fragments, but contain only a few 3D domains—or even just one. Crucially, only 3D domains are relevant for *in vitro* experiments that require isolating parts of the macromolecule capable of folding or functioning on their own.

In the particular case of ribosomal RNAs, for over forty years, authors have partitioned their structures into well-established domains that are evolutionarily conserved [27–29]. Although comparable in size to 3D domains, these substructures originate from functional and comparative studies across organisms, and are based on 2D structural organization rather than 3D geometry. This raises the question of how closely the RNA 3D domains identified by our segmentation method align with those based alternative definitions. In the case of proteins, it has been shown that domain annotations from the CATH, SCOP, and ECOD databases often diverge, reflecting the different criteria used to partition 3D structures [30]. In this work, we addressed this issue by incorporating domain annotations from the Rfam database [31] into our benchmark. The goal was to evaluate the relevance of the domains delimited by RNA3DClust in the light of those defined in Rfam, which are based on sequence conservation, secondary structure, and known biological functions (see Section 2.5.3).

### 2.2. Candidate clustering algorithms

To delimit 3D domains within RNA structures, a clustering algorithm should meet the following three requirements:

● As the length of an RNA chain can reach tens of thousands of residues, it is expected to contain folded regions, which may be separated by unfolded RNA segments: the above-defined linkers. The atoms of these linkers should not be assigned to any specific cluster. Therefore, the clustering algorithm should be robust against outliers, *i.e.* it should explicitly identify orphans, which are the coordinate points that do not belong to any meaningful cluster in the RNA structure.
● As the physicochemical forces involved in the process of RNA folding differ from those that operate in proteins, the polynucleotide chain may assume a wider variety of shapes that are not necessarily globular. Therefore, the clustering algorithm should be able to find clusters of irregular shapes and sizes.
● As the number of 3D domains is expected to vary from one RNA conformation to another, the clustering algorithm should be “non-parametric”, *i.e.* it should not require the number of clusters in advance.

We looked for these properties among commonly-used clustering algorithms (Table 1): *k*-means [32], hierarchical clustering [33], density-based spatial clustering of applications with noise (DBSCAN) [34], hierarchical DBSCAN (HDBSCAN) [35], Mean Shift [18], Gaussian mixture models (GMM) [36], spectral clustering [37], self-organizing maps (SOM) [38], and the highly connected subgraphs (HCS) clustering algorithm [39]. It appeared that the above three criteria only match with the definition of density-based clustering algorithms, which group data points based on the density of their neighborhoods. In the present comparison, these are the DBSCAN, HDBSCAN, and Mean Shift algorithms.

**Table 1.** Comparison between widely-used clustering algorithms, based on the three criteria that we identified as necessary for segmenting RNA 3D structures. The “*” symbol denotes that the criterion is met only under certain conditions (see Section 2.2.2 for details).

| Algorithm | Robust against outliers | Find irregular shapes | Non-parametric |
| --- | --- | --- | --- |
| DBSCAN | ✓ | ✓ | ✓ |
| GMM |  |  |  |
| HCS |  | ✓ | ✓ |
| HDBSCAN | ✓ | ✓ | ✓ |
| Hierarchical |  | ✓ | * |
| $k$-means | | | |
| Mean Shift | ✓ | ✓ | ✓ |
| SOM |  | ✓ |  |
| Spectral | * | ✓ | * |

#### 2.2.1. Selected algorithms

The **DBSCAN** algorithm has two main hyperparameters: the neighborhood radius *ε* and minimum neighbors (MinPts). The hyperparameter *ε* defines the maximum distance for points to be considered neighbors, while MinPts indicates the minimum number of neighbors a point needs to be considered a cluster “core”. DBSCAN begins by randomly selecting a not-yet-visited point in the dataset, and if it has at least MinPts neighbors within a sphere of radius *ε* centered on it, then it is considered as a core point. Every time a core point is found, the clustering algorithm is initiated with that point. Once a core point is identified, DBSCAN expands the cluster by looking at all points within *ε* distance from this core point. If any of these neighboring points are also core points, their neighbors are included into the cluster. This process continues, adding all reachable points within *ε* distance until no new point can be added to the cluster. If a point is not a core point and is not reachable from any other core point, it is classified as outlier, *i.e.* not belonging to any cluster. For DBSCAN to capture meaningful clusters effectively, choosing the right values for *ε* and MinPts is essential.

**HDBSCAN** extends DBSCAN by converting it into a hierarchical clustering algorithm and then extracting the most stable clusters. It eliminates the need to choose *ε*, DBSCAN’s most sensitive hyperparameter, by considering all possible density thresholds and constructing a hierarchy of clusters. Specifically, HDBSCAN transforms the space by computing mutual reachability distances instead of Euclidean distances, where these distances are based on the *k*-th nearest neighbor and used to define local density. Although the hyperparameter *k* is conceptually similar to MinPts in DBSCAN, it is used differently: in HDBSCAN, *k* defines the neighborhood size for estimating core distances, which influence mutual reachability, rather than directly determining core points or cluster membership. A minimum spanning tree (MST) is then built from the mutual reachability graph, and the hierarchy is constructed by progressively removing the weakest edges in the MST. The resulting tree is condensed using a hyperparameter denoted “mpts”, which defines the minimum number of points a cluster must contain to be considered valid. A stability analysis is then applied to identify and retain the most persistent clusters in the hierarchy. Finally, a flat clustering is extracted by selecting the most stable clusters at different levels of the tree, although the full hierarchy can also be visualized.

More recently, a **hybrid approach** [40] combining DBSCAN and HDBSCAN has been proposed to mitigate over-segmentation into micro-clusters in dense regions when using small values for the mpts (minimum cluster size) hyperparameter. This method introduces an *ε*-like distance threshold that constrains how far apart points can be within a flat cluster. In the original formulation, this threshold is applied during the construction of the condensed cluster tree, where it effectively prevents the split of cluster components that are too close in mutual reachability distance. In contrast, some implementations (like in scikit-learn [41]) apply this threshold after the hierarchy has been constructed, during the flat cluster selection step, where it serves as a merging criterion for adjacent cluster components. Overall, this hybrid approach preserves HDBSCAN’s ability to detect clusters across varying densities, while introducing control over cluster compactness. For this reason, we selected it as one of the candidate clustering algorithms in our study.

The **Mean Shift** clustering identifies dense regions in data by iteratively shifting each data point towards the “center of mass” of its neighbors. The algorithm uses a kernel function, essentially a weighting function, to determine the influence of each neighbor on the shift direction. Mean Shift relies on two hyperparameters. The first one is the kernel type, which defines how the influence of neighbors is distributed. A uniform kernel gives equal weight to all neighbors within the search radius. Other types, like Gaussian kernels, assign higher weights to closer neighbors, creating a smoother shift towards the center of mass. The other hyperparameter is the bandwidth (or window size) *h*, which defines the search radius around each data point. A larger size considers more neighbors, potentially smoothing out smaller clusters and merging them into larger ones. Conversely, a smaller size focuses on closer neighbors, potentially identifying smaller, more distinct clusters. By repeatedly shifting points toward their local density peaks based on the kernel and its neighbors, the algorithm converges to the centers of clusters.

#### 2.2.2. Excluded algorithms

Among the algorithms that do not satisfy all three conditions for clustering RNA 3D structures (Table 1), both GMMs and *k*-means fail to meet any criterion. GMMs assume clusters follow a Gaussian distribution, resulting in elliptical cluster boundaries in the original feature space, while *k*-means assumes spherical, equally sized clusters. In both cases, these shape assumptions make them poorly suited for non-convex or highly irregular cluster structures. Although the number of clusters *k* can be estimated for GMMs using model selection criteria such as BIC or AIC, and for *k*-means using methods such as the silhouette score or gap statistic, these approaches can be computationally costly and are often unreliable when the underlying cluster geometry deviates strongly from the model assumptions.

Both HCS and hierarchical clustering fall short due to their handling of outliers. In HCS, outliers are not explicitly modeled: all nodes are assigned to some cluster unless a manual or heuristic post-processing step is applied to prune weakly connected nodes. Similarly, hierarchical clustering does not have a built-in mechanism for distinguishing noise or outliers from legitimate data points. As a result, outliers can significantly influence the dendrogram structure, especially with linkage methods like single or complete linkage. While hierarchical clustering does not require predefining the number of clusters, selecting a cutoff to extract flat clusters from the dendrogram can be subjective and requires exploratory interpretation.

In the case of SOMs, while they preserve topological relationships in the input data, the resulting grid is not inherently clustered. Final clustering typically requires post hoc methods (e.g., k-means or U-matrix thresholding), which may not accurately capture irregular cluster shapes or densities. Moreover, SOMs are sensitive to outliers, as such points can disproportionately influence the neighborhood updating during training. Overall, SOMs are better understood as dimensionality reduction and visualization techniques rather than standalone clustering algorithms.

Although spectral clustering is well-suited for detecting clusters with non-convex or irregular shapes, it is generally not robust to outliers. This is because it tends to assign all points, including noise, to one of the clusters, often based on weak or spurious affinities. Whether spectral clustering requires specifying the number of clusters in advance depends on the method used after spectral embedding. Typically, *k*-means is applied in the embedded space, which does require a predefined *k*, but alternative strategies exist: the eigengap heuristic can provide a direct estimate of *k* [42], while model selection approaches such as the silhouette score evaluate clustering quality across different candidate values of *k* [43]. However, in this study, we restrict our attention to algorithms that inherently determine the number of clusters, without relying on such strategies.

### 2.3. Post-clustering procedure

As the selected clustering algorithms only rely on the density of points, the output clusters may be scattered along the RNA sequence, which is not biologically meaningful. To tackle this critical problem, we have developed a post-clustering procedure, which consists of two main steps: processing outliers and processing labeled clusters. For that purpose, we have defined eight rules for the post-clustering procedure (Fig. 2). These rules are described subsequently, in their order of application:

● Process of outlier clusters: the outliers located inside another cluster are relabeled according to that cluster (Fig. 2A); the outliers with length < 10 nt located between two clusters are labeled half as the cluster on the left and half as the cluster on the right (Fig. 2B); the outliers with length < 10 nt or ≥ 100 nt located on the right end of the molecule and besides a cluster with length ≥ 30 nt are labeled according to that cluster (Fig. 2C). The same process is applied to the outliers located on the left end. If a labeled cluster is < 30 nt and located between two outlier clusters that both have a length ≥ 30 nt, then they become a unified outlier segment (Fig. 2D).
● Process of labeled clusters: if the labeled cluster has a length < 30 nt and is located in the middle of a larger cluster whose length is > 30 nt, then it is relabeled according to the larger cluster (Fig. 2E); if the labeled cluster whose length is < 30 nt is located on the right end of the molecule and on the left is a larger labeled cluster whose length is > 30 nt, then it is relabeled again according to the larger cluster (Fig. 2F). The same process is applied to the labeled cluster on the left end; if the labeled cluster has a length < 30 nt and is located between two larger clusters whose length are > 30 nt, then it is labeled half according to the cluster on the left, half according to the cluster on the right (Fig. 2G); if the labeled cluster has a length > 30 nt and is located between a larger cluster whose length is > 30 nt and an outlier region, then it is relabeled according to the label cluster (Fig. 2H).

**Figure 2.**
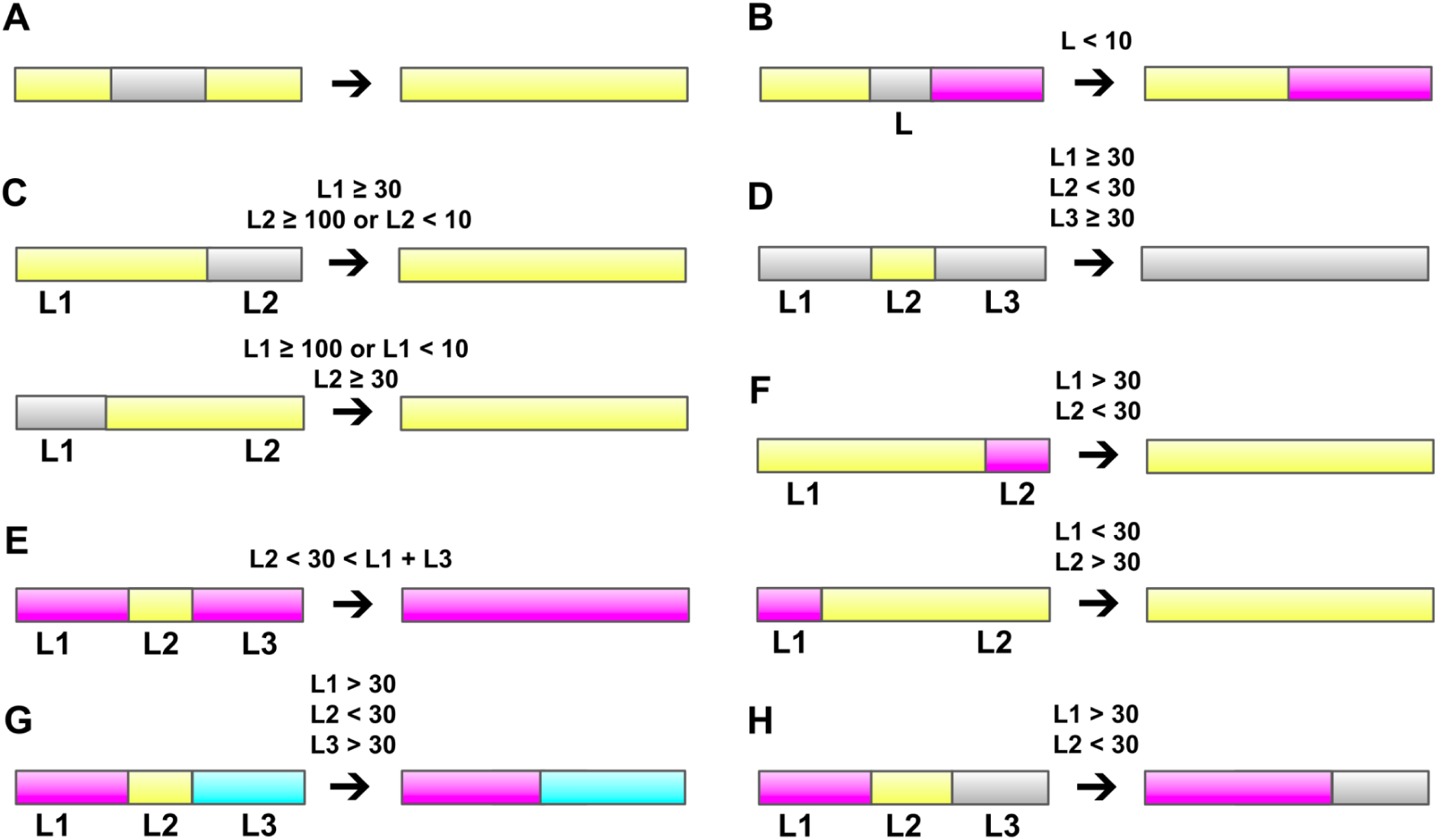
The eight rules for the post-clustering procedure. (**A**, **B**, **C**, and **D**) The rules for deleting or extending outliers; (**E**, **F**, **G**, and **H**) The rules for labeled clusters. The gray color indicates outlier regions. The yellow, magenta and cyan colors indicate cluster regions. The sequence length of the left, middle and right regions are symbolized by “L1”, “L2” and “L3”, respectively.

The above two processes do not create new clusters, but only expand or remove clusters. Since we want to avoid cluster fragmentation, expanding clusters is preferred over cluster deletion. Therefore, processing the outlier cluster (narrowing the outlier cluster) is carried out first, followed by processing the label cluster (narrowing the label cluster). The above two processes are repeated until the result is unchanged.

### 2.4. Segmentation quality scores

To assess the similarity between the domain decomposition produced by RNA3DClust and the ground-truth reference, we used three metrics. The first two are the most cited scores in the field of protein domain parsing, the normalized domain overlap (NDO) [44] and the domain boundary distance (DBD) [45], which we managed to directly apply to RNA. The third metric has been developed for this work as a compromise between the NDO and DBD scores, and has been named chain segment distance (CSD). Finally, as a complement to these metrics, we evaluated the correctness of the decomposition in terms of the number of delineated domains. For that purpose, we measured the performance of RNA3DClust at classifying the input RNA as being either a single- or multi-domain structure, similarly to previous studies about protein 3D domains [20,46,47]. Thus, the resulting classification performance, as well as the NDO and DBD scores, can be placed side by side with those of state-of-the-art protein domain parsers. However, the latter can only serve as a point of reference, as no direct comparison is possible between RNA and protein molecules.

#### 2.4.1. Normalized domain overlap (NDO)

To assess the correctness of a 3D domain parser, the sequence overlap between the computed domains and their ground-truth counterparts is measured. As simply comparing the domain boundaries may introduce biases, authors have developed a formula called normalized domain overlap (NDO) [44]. Here, “overlap” designates the number of common residue indices between two domain annotations. First, for each domain *i*, a net overlap score is calculated as:

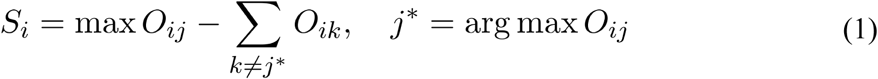

where max *O_ij_* selects the highest overlap for domain *i* among all possible domains *j*, and ∑*_k≠j_*_*_ *O_ik_*subtracts all other overlaps, to remove over-segmentations. Domain *j*\* is the one that has the highest overlap with domain *i*, excluding linkers. Domain *k* is any other overlapping domain, including linkers. The net overlap scores *S_i_* are calculated for all non-linker domains *i*, from both the computed and ground-truth domain decompositions. Then, normalized overlap score is calculated as:

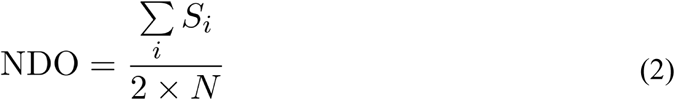

where the sum of scores *S_i_* is averaged for both the computed and true domains, and normalized by the total number of true domain residues *N*, excluding linkers and missing residues. The NDO was designed with values ranging from 0.0 to 1.0, and the higher the NDO the better the quality of the domain decomposition, *i.e.* high overlap and minimal over-segmentation. In practical applications, however, negative values can occur—although rarely—and indicate either a significant over-segmentation (many small computed domains overlapping the same true domain), or under-segmentation, or a severe misalignment between the computed and true domains.

#### 2.4.2. Domain boundary distance (DBD)

The domain decomposition results were also assessed using the DBD score [45], which is based on the distances between the computed and ground-truth domain boundaries, denoted by the letters *i* and *j*, respectively. Each computed boundary *i* within *T* residues of a true boundary *j* is evaluated with the score:

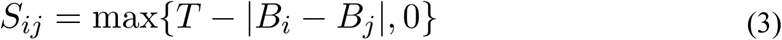

where *B_i_* is the computed boundary position (residue index), *B_j_* is the true boundary position, and ∣*B_i_*− *B_j_*∣ is the absolute difference (or distance) between them. Here, the boundary position between two domains is a single number representing the 3’-end terminal nucleotide of the upstream domain. If the difference ∣*B_i_*− *B_j_*∣ is greater or equal to the threshold *T*, the score becomes 0 (no contribution). Importantly, due to the threshold *T* and the minimum sequence length of a domain (Section 2.1), the score *S_ij_*of a given computed boundary *i* can be > 0 only for a single true boundary *j* (*i.e.* the closest one). Then, the DBD is calculated as the total score for the domain decomposition:

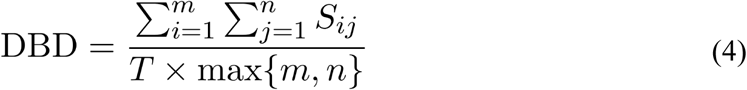

where *m* and *n* represent the number of domain boundaries in the computed and true decompositions, respectively. The value of the DBD ranges from 0.0 to 1.0, and the higher the DBD the better the quality of the domain decomposition, *i.e.* the more accurate the computed boundaries. Importantly, if the domain boundary has a linker, the whole linker is regarded as the domain boundary. For the threshold *T*, different values have been used in the literature, either *T* = 8 residues [19,45], or *T* = 20 residues [46,48]. If this margin *T* is too narrow to cover the deviation ∣*B_i_*− *B_j_*∣, the score in Eq. 4 will be 0, which may underrate the quality of the computed boundary. Although this is not a problem for highly accurate methods, RNA3DClust is not expected to match the accuracy of its well-established protein-specific counterparts, as it is the first 3D domain parser dedicated to RNA. We therefore used the less strict margin of *T* = 20 residues.

#### 2.4.3. Chain segment distance (CSD)

Although the NDO and DBD scores are often used as complementary metrics, they each have drawbacks. In the case of a segmentation with an incorrect number of domains, the NDO value may still be high, due to the number of overlapping residues. In the case of a segmentation where the boundaries are only slightly off, the DBD value may still be low, due to the zeros in the numerator (Eq. 4), which result from the use of a too small threshold (Eq. 3). In addition, when the DBD was released, most of the available domain parsers were not able to detect linkers. Therefore, this metric was designed to compare domain decompositions in which only the ground truth contains linker regions. This latter shortcoming, along with the fact that the DBD is not suited for single-domain annotations (*i.e.*, no segmentation), may lead to score values that may not properly reflect the segmentation quality.

Here, we present a new scoring function aimed at offering a compromise between the NDO and DBD. Like the DBD, the scoring follows a decreasing pattern based on a distance threshold. However, rather than just boundaries, this distance is calculated between domains, like the NDO. For each computed domain *i* and true domain *j*, a score *S_ij_*is calculated as:

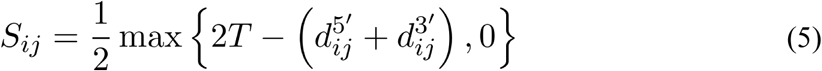

where *d_ij_*^5’^ is the distance (in nt) between the 5’ end of the domain *i* and that of the domain *j*. Similarly, *d_ij_*^3’^ is the distance between the 3’ ends. The threshold *T* serves the same purpose as in the DBD formula (Eq. 3) and was also set to *T* = 20 residues. Regarding linkers, their length is added to that of the adjacent domain, only if it lowers the value of *d_ij_*^5’^ + *d_ij_*^3’^; otherwise, the length of the linker is not counted. Finally, the chain segment distance (CSD) score is calculated as a piecewise function:

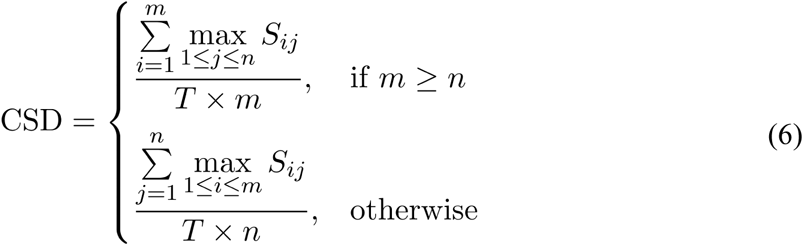

where *m* and *n* are the total numbers of computed and true domains, respectively. The value of the CSD ranges from 0.0 to 1.0, and the higher it is, the better the quality of the segmentation. Finally, it should be noted that the chosen name for this new score conveys the idea of a usage not restricted to RNAs, nor domains—although, it is the object of the present work. The CSD could be applied to protein chains, as well as other structural elements, such as subdomains or supersecondary structures.

#### 2.4.4. Single- vs multi-domain classification

The fourth performance assessment evaluates RNA3DClust as a binary classifier that distinguishes between single- and multi-domain 3D structures. As “single-domain” actually designates 3D structures that cannot be separated into different substructures, it is the ability to detect segmentable structures that is tested here. Such evaluation of 3D domain parsers has been used multiple times in the protein-related literature [20,46,47]. The RNA3DHub, and RNA3DB sets are balanced between the two classes (single- and multi-domain), whereas the LNCipedia set is not. Therefore, performance metrics chosen were the accuracy (Acc) and the Matthews correlation coefficient (MCC), calculated as:

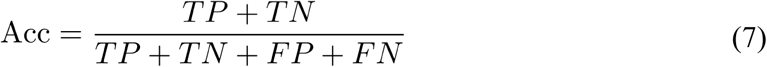

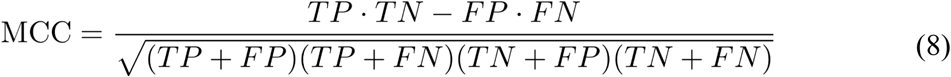

where the true positives (TP) and false positives (FP) are the ground-truth multi-domain structures correctly and incorrectly detected (*i.e.*, based on domain counts), respectively; similarly, the true negatives (TN) and false negatives (FN) are the ground-truth single-domain structures correctly and incorrectly detected by RNA3DClust, respectively.

### 2.5. Creation of the datasets

As the concept of RNA 3D domains has never been developed in the literature, we had to define datasets by (i) selecting representative RNA conformations, and (ii) manually annotating domain boundaries based on Wetlaufer’s visual criteria: compactness and spatial separation [17]. Thus, we created the following non-redundant datasets, named after the web resources from which they were extracted (details are provided in the subsections below):

● The “RNA3DHub” (*n*=132 RNA chains) and “RNA3DB” (*n*=26 RNA chains) datasets are both made of experimentally determined 3D structures, and were both used for the hyperparameter tuning and cross-validation procedures (Tables S1 and S2, respectively).
● The “LNCipedia” (*n*=69 RNA chains) dataset contains exclusively predicted 3D structures of lncRNAs, and was used for benchmarking purposes (Table S3).
● The dataset “Rfam consensus” (*n*=122 domains) represents a complementary ground-truth to the aforementioned Wetlaufer’s visual criteria (Table S4). It is made of annotations that are in agreement between the evolutionary and functional ones from

the Rfam database, and those based on 3D geometrical criteria. Importantly, each entry in this dataset is a domain rather than an RNA chain, as multiple Rfam domains may overlap within a single RNA chain.

#### 2.5.1. RNA3DHub and RNA3DB sets

For the hyperparameter tuning and cross-validation procedures, we used a representative set of RNA 3D structures as defined by the RNA 3D Hub web resource [49]. From the release 3.390, we took only the structures with a crystallographic resolution ≤ 2.5 Å and an RNA sequence length ≥ 100 nt. This latter threshold represents the minimum length necessary to accommodate two 3D domains and their linker, with only nucleotides having coordinates in the PDB file being counted. Thus, our “RNA3DHub” set contains 132 non-redundant RNA chains, corresponding to 66 PDB entries. These belong to a variety of RNA types (such as ribosomal RNAs, riboswitches, or ribozymes) and to organisms across all kingdoms of life and viruses. The RNA sequence lengths range from 100 nt to 3764 nt, with a mean of 1049,7 nt and a median of 190 nt. We assigned the 3D domain delimitations by visual inspection, using the PyMOL Molecular Graphics System, Version 2.5, Schrödinger, LLC. We thus obtained a total of 82 single-domain RNAs and 50 multi-domain RNAs (46 two-domain RNAs and 4 three-domain RNAs).

Recently, authors have proposed a new approach to tackle the problem of redundancy when devising datasets of RNA 3D structures, with the publication of the RNA3DB dataset [50]. Thus, to assess how the obtained hyperparameter values depend on the set RNAs, we used RNA3DB as an alternative to RNA 3D Hub. Out of the 127 distinct groups (called “Components”) of the RNA3DB set from December 17, 2024, we selected the longest representative RNA chain of each, and then filtered those having a resolution ≤ 3.5 Å and an RNA sequence length ≥ 100 nt. This resulted in a total of 26 structurally-dissimilar RNA chains, corresponding to 26 PDB entries, with the shortest and longest being 100 nt and 4269 nt, respectively. After our manual annotation of 3D domains, we obtained a total of 16 single-domain RNAs, 9 two-domain RNAs, and a single three-domain RNA.

#### 2.5.2. LNCipedia set

The subject of lncRNAs has steadily gained attention over the recent years, with more and more of these long transcripts being found in RNA-seq experiments related to various fields of biology, such as human pathologies or plant physiology. Due to their sequence length ranging from 200 nt to thousands of residues, lncRNAs are likely to contain structured and compact regions, separated by stretches of unfolded nucleotide chain. Unfortunately, no experimentally determined structure of a complete lncRNA is currently available in the PDB. Therefore, to assess the capacity of RNA3DClust to annotate domains in such cases, we generated theoretical 3D structures of lncRNAs. For that purpose, we took ncRNA sequences from the human lncRNAs database LNCipedia [51], then used AlphaFold3 [5] for predicting their 3D structures. Thus, our LNCipedia set contains structural predictions of 69 lncRNAs (15 single- and 54 multi-domain structures) and is independent from the RNA3DHub and RNA3DB sets regarding the content in RNA 3D structures.

In all attempts to generate 3D structural models of lncRNAs using AlphaFold 3, the predicted TM-score (pTM) never exceeded 0.25, with all values falling in the range 0.08–0.24 (Table S3). Such values would typically indicate low confidence in the global fold when interpreting AlphaFold predictions for proteins. However, it is important to note that AlphaFold’s confidence metrics, including pTM, are calibrated on protein structures and a limited dataset of RNAs available in the PDB, from which lncRNAs are essentially absent. Consequently, the low pTM values observed here likely reflect the model’s lack of prior knowledge for this class of RNAs rather than necessarily indicating that the predicted structures are inaccurate. In addition to pTM, we report the per-nucleotide confidence metric pLDDT (predicted Local Distance Difference Test), which provides a local measure of structural reliability. For the same reason as with the pTM scores, the pLDDT values must be interpreted carefully: they are best used relatively, to compare confidence between regions within the same model, rather than relying on absolute cutoffs. The pLDDT values for each 3D model are provided in JSON files (project repository).

#### 2.5.3. Rfam consensus set

Manual annotation of 3D domains has a long history in structural biology, exemplified by decades of expert-curated entries in the SCOP database. The “Islam dataset” of protein 3D domains, used for benchmarking domain decomposition methods, is another example of manually defined annotations. In the current versions of the SCOP and CATH databases, the addition of new 3D domain annotations is only semi-automated. More generally, in machine learning, “ground truth” labels, such as those used in image segmentation, are often established by human annotators. However, manual annotation is inherently subjective, and the robustness of the resulting dataset depends on both the expertise and the number of annotators involved.

Therefore, to minimize subjectivity, we chose to evaluate our method against a ground truth that is externally-defined and different from our 3D-based annotations. Specifically, we used RNA domains from the Rfam database, whose definitions are based on sequence conservation, secondary structure, and biological function. From a technical standpoint, we used Infernal [52] to query Rfam entries for each structure in the RNA3DHub set. From the structures of these RNA chains, we extracted the nucleic acid sequences, and then ran Infernal on them to get the corresponding Rfam entries. Thus, out of the 132 RNA chains of the RNA3DHub set, 117 had one or several Rfam domains, for a total of 272 Rfam domains.

The criteria used in Rfam differ fundamentally from those used to delimit RNA 3D domains, and therefore one can reasonably expect discrepancies between the two annotation schemes. For proteins, such differences have been documented: domain annotations from the CATH and SCOP databases agree in only ∼78% of cases [30]. Following the approach of the “Consensus set” defined in that previous study, we here identify a subset of Rfam annotations that agree with our RNA 3D domain boundaries. For each Rfam domain, agreement was quantified using the Jaccard index—also known as the intersection-over-union (IoU)—a metric recently applied in the protein 3D domain literature [53]. Thus, for a Rfam domain *D*_i_ ^Rfam^, the IoU is calculated with its corresponding 3D domain *D*_j_ ^3D^, which is the one that shares the largest intersection (*i.e.* overlap) among all of the *n* RNA 3D domains:

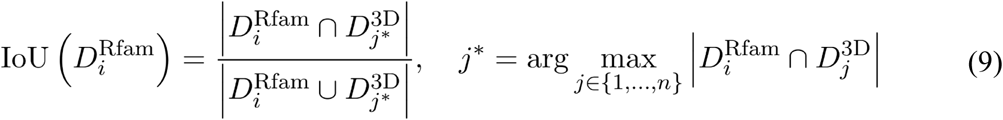

In accordance with the literature, we used a minimum IoU value of 0.8 for considering two annotations to be in agreement [53]. Thus, out of the 272 Rfam domains identified in the RNA3DHub dataset, 122 showed agreement with our 3D domain boundaries, corresponding to an agreement rate of ∼45%. We followed the same approach to define a consensus set between Rfam domains and our visual annotations of the 3D structures from RNA3DB. However, out of the 16 entries, only 6 had an IoU ≥ 0.8, preventing the constitution of a second consensus dataset—we still included some of these entries as examples in the Section 3.4 below and in Table S5.

To evaluate the performance of RNA3DClust on this Rfam consensus set, we also calculated the IoU as in Eq. 9, using the same 0.8 threshold, but with *D*_j_ ^3D^ representing computed, rather than visually-delimited, 3D domains. We used the IoU instead of the segmentation quality scores described in Section 2.4, due to a peculiarity of Rfam domain annotations: unlike 3D domains, Rfam annotations within a single RNA molecule may overlap, making segmentation metrics inapplicable. Finally, it should be noted that the 181 cases of disagreement were not analyzed further in this work. Addressing such a “dissensus set” would require an algorithm capable of producing multiple alternative domain decompositions for a given RNA, analogous to the SWORD approach used for proteins [30].

### 2.6. Statistical analysis

#### 2.6.1. Tests for significance

To compare the different average values of the NDO, DBD, and CSD scores, we used the Student’s t-test and the Wilcoxon signed-rank test, for normally and non-normally distributed data, respectively, with a *p*-value threshold of 0.05. To assess the normality of the distributions, we used the Shapiro–Wilk test, with a *p*-value threshold of 0.05. To assess the linear relationship between these scores, we calculated the Pearson (*ρ*) correlation coefficient. The *p*-value quantifies how likely it is to observe a correlation coefficient as extreme as the one calculated, assuming the null hypothesis is true—that is, assuming there is no actual correlation in the population (*ρ*=0). All these statistics and related plots were computed with the Python 3.10 modules SciPy and Matplotlib.

#### 2.6.2. Cross-validation

The RNA3DHub set (and, alternatively, the RNA3DB set) was used for both hyperparameter tuning and benchmarking purposes, which might introduce an obvious bias. Therefore, to assess any potential overestimation of the performance, the score values calculated on the two-domain RNAs were compared to those obtained by following a group *k*-fold cross-validation procedure. The latter was defined by splitting the subset of two-domain RNAs into *k* groups, based on the RNA types defined in RNA 3D Hub or RNA3DB. Then, for each of these two-domain structures, the scores were calculated using hyperparameters that were tuned on the *k*−1 groups it does not belong to. For example, the scores of the 11 18S rRNAs of the RNA3DHub set were calculated using hyperparameters tuned on the 121 other RNA structures. Finally, the reason for excluding single-domain RNAs from the hyperparameter tuning and cross-validation is that it would otherwise bias the results, since the propensity of Mean Shift to generate a single cluster increases as the window size (or bandwidth) grows.

#### 2.6.3. Bootstrap resampling

To determine whether the difference between the MCC values calculated for the RNA3DHub, RNA3DB, and LNCipedia sets is statistically significant, we performed a bootstrap-based test. This was done by (i) resampling with replacement from the original data within each dataset separately, then (ii) recalculating the MCC for each resampled dataset, and (iii) computing the MCC difference between the two datasets. The resampled datasets were of the same size as the original datasets: 132, 26, and 69 entries, for the RNA3DHub, RNA3DB, and LNCipedia sets, respectively. These three steps were repeated 10,000 times to generate an empirically estimated distribution of the MCC differences. Finally, the *p*-value was computed by checking how extreme the observed difference is compared to that null distribution. For example, a *p*-value of 0.05 means that, out of the 10,000 MCC differences generated, only 500 were greater than the observed difference. It should be noted that this bootstrapping was chosen over a permutation test, due to the non-exchangeability of the data between the RNA3DHub, RNA3DB, and LNCipedia sets (*i.e.*, experimental vs predicted structures).

## 3. RESULTS AND DISCUSSION

### 3.1. Initial feasibility assessment

In this section, we describe how we assessed the feasibility of exploring the hyperparameter space of the Mean Shift, DBSCAN, and HDBSCAN clustering algorithms, based on the 132 structures in our RNA3DHub set of annotated RNA 3D domains.

For DBSCAN, we have performed the preliminary tests by discretizing the bidimensional hyperparameter space, for both *ε* and MinPts (Figs. 3A and 3B). The two hyperparameters were expressed in ratio, with discretization step sizes of 0.1: the neighborhood radius *ε* was calculated based on the ratio of the search radius to the maximum distance between two points; MinPts was calculated based on the ratio of the number of neighbor points to the number of total points. Thus, we tested a combination of 10 values (from 0.1 to 1.0) of *ε* and of MinPts, for a total of 100 pairs of hyperparameter values. For the subset of 82 single-domain structures, several combinations of values resulted in more than 50% of correct annotations (magenta dots in Fig. 3A). However, for the subset of 50 multi-domain structures, we observed no pair of hyperparameter values that can return the correct number of RNA 3D domains (white dots in Fig. 3B). This means that finding a pair of DBSCAN hyperparameter values that results in the correct number of domains for the entire RNA3DHub set—if such a pair of values exists—would require an even smaller discretization step size and, therefore, an exploration process that could be unaffordably long. For this reason, we do not use the DBSCAN algorithm in the rest of the article.

**Figure 3.**
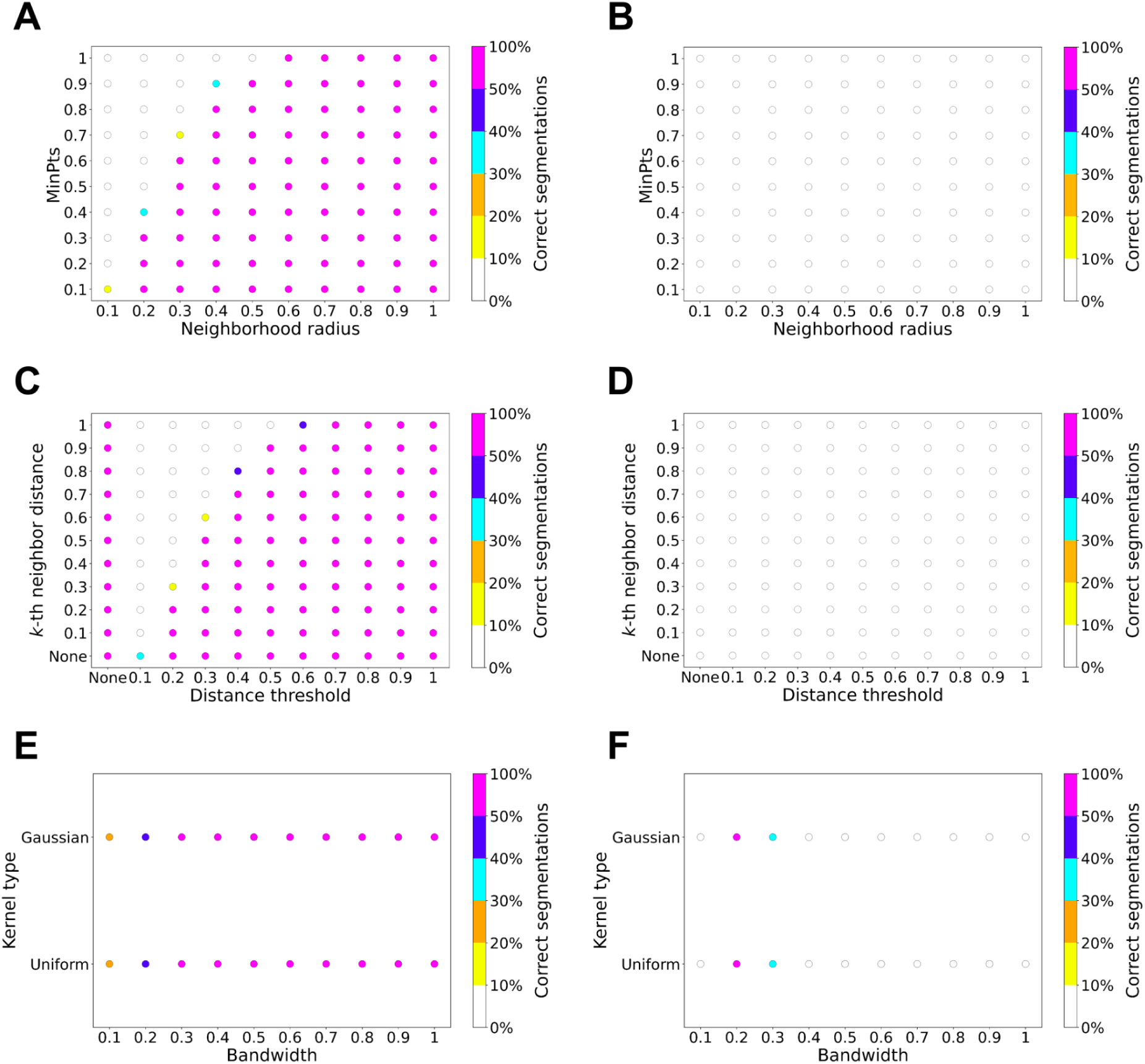
Scatter plots representing the search for hyperparameter values that return a correct segmentation in terms of number of domains, for (**A**, **B**) DBSCAN, (**C**, **D**) HDBSCAN and the hybrid approach, and (**E**, **F**) Mean Shift. The computations were performed on the 132 experimental RNAs structures from the RNA3DHub set: (**Left panels**) The subset of 82 single-domain annotations; (**Right panels**) The subset of 50 multi-domain annotations. The color of each dot represents the percentage of cases with the right number of clusters, as shown by the color scale.

A similar hyperparameter exploration was also conducted for HDBSCAN, employing the same ratio calculation and discretization step size as used for DBSCAN (Figs. 3C and 3D).

Unlike DBSCAN, HDBSCAN does not require an *ε* parameter; however, a subsequently proposed hybrid approach introduced an *ε*-like distance threshold as an additional hyperparameter to address over-segmentation (see Section 2.2.1). Thus, on the *x*-axis, a value of “None” for the distance threshold denotes the original HDBSCAN algorithm, whereas the other values (from 0.1 to 1.0) correspond to the hybrid approach. The *y*-axis represents the discretized ratio value of HDBSCAN’s *k* hyperparameter. When *k* is set to “None”, it defaults to the mpts hyperparameter, which defines the minimum number of points required for a cluster to be considered valid (see Section 2.2.1). In the present work, mpts was fixed at 30 nt, corresponding to the minimum length of an RNA 3D domain. Thus, 11 combinations of hyperparameter values were tested for HDBSCAN (distance threshold = “None”, with *k* ranging from “None” to 1.0), and 110 combinations for the hybrid approach. As with DBSCAN, correct segmentations were observed exclusively for the subset of single-domain annotations (Fig. 3C), whereas the correct number of domains was never found for the other subset (Fig. 3D). Consequently, neither HDBSCAN nor the hybrid approach were considered further in subsequent investigations.

Finally, the hyperparameter space of Mean Shift was also explored (Figs. 3E and 3F), using 2 different kernel types (Gaussian and uniform) and 10 bandwidths, proportional to the quantiles of the original data distribution (with values ranging from 0.1 to 1.0, in intervals of 0.1). To add the possibility of using a Gaussian kernel, we modified the source code of the scikit-learn implementation of Mean Shift [41]. We kept the default convergence threshold: if all points move by less than 0.001 Å between iterations, the algorithm assumes convergence and stops. The results on the single-domain annotations in the RNA3DHub set (Fig. 3E) were computed solely for the sake of completeness, but were not taken into account, as Mean Shift is biased toward this subset. Indeed, with the Mean Shift algorithm, using wider bandwidths yields fewer and larger clusters. Thus, a very large bandwidth would systematically generate single clusters, which would be counted as correct for all the entries annotated as single-domain RNA. For the subset of multi-domain annotations (Fig. 3F), out of the 20 pairs of hyperparameter values, we observed that a 0.2 bandwidth with either type of kernel resulted in the correct number of clusters, for more than 50% of the cases. This indicates that there may exist a pair of hyperparameter values that allow to correctly delimit 3D domains for the RNAs of the RNA3DHub set. Therefore, we decided to further tune the two hyperparameters of Mean Shift: the kernel type and the bandwidth.

In addition to the RNA3DHub set, this initial feasibility assessment was also performed on the smaller RNA3DB set (Fig. S1), where Mean Shift again emerged as the only viable method for segmenting RNA 3D structures. Besides, across the three single-atom representations tested (C1’, C3’ and C4’), the results differed only marginally, such that their respective plots correspond to that shown in Figure 3 (due to the discretized % of correct segmentations), thereby leading to the same conclusions regarding the clustering algorithms. Nevertheless, we carried out additional comparative tests on the choice of atom in the following sections. Finally, despite Mean Shift predating DBSCAN, HDBSCAN, and the hybrid method, the overall conclusion of this feasibility assessment is not unexpected, as Mean Shift continues to be relevant in contemporary literature. For example, GPU-accelerated implementations have recently been proposed for image segmentation and object tracking [54], and recent research has integrated Mean Shift into contrastive representation learning frameworks [55].

### 3.2. Hyperparameter tuning

To determine the optimal bandwidth and kernel type for Mean Shift, we performed hyperparameter tuning based on the outputs of the clustering algorithm and the subsequent post-processing described above (Fig. 2). Although the Mean Shift algorithm itself is unsupervised, the use of labeled data (*i.e.*, datasets of 3D domain annotations) for hyperparameter optimization qualifies the overall pipeline as semi-supervised. In contrast to the feasibility assessment in the previous section, segmentation quality was evaluated here using three scoring functions: NDO, DBD, and the newly developed CSD (Eqs. 5 and 6). The values presented correspond to the C3’-based segmentation (results for C1’ and C4’ are provided in the Supplementary Materials). Hyperparameter tuning was carried out on the subset of multi-domain structures from the RNA3DHub set, due to the aforementioned bias of Mean Shift toward single-domain annotations. For this subset, we can observe in Fig. 4 that the bandwidth affects the result more than the kernel type. Specifically, there is no significant difference between the uniform and the Gaussian kernel types, for the same bandwidth value. This was not unexpected since, when using a fixed bandwidth, the kernel shape only matters when the data is very sparse or has extreme outliers. We also found that a uniform kernel with a bandwidth of 0.2 quantile yields the highest average NDO (0.744), as well as the highest average DBD (0.348) and CSD (0.557). The average NDO calculated by cross-validation was identical (0.744), which means that these performance results are not overestimated. We therefore set “bandwidth = 0.2” and “kernel = uniform” as default for the Mean Shift clustering procedure in the RNA3DClust tool, and used these settings for the rest of the article. This choice was further supported by hyperparameter tuning on the RNA3DB dataset, which yielded the same settings (Fig. S2). Finally, we observed that the optimal parameter values were identical across the three single-atom representations tested (C1’, C3’, and C4’; Figs. S2 and S3).

**Figure 4.**
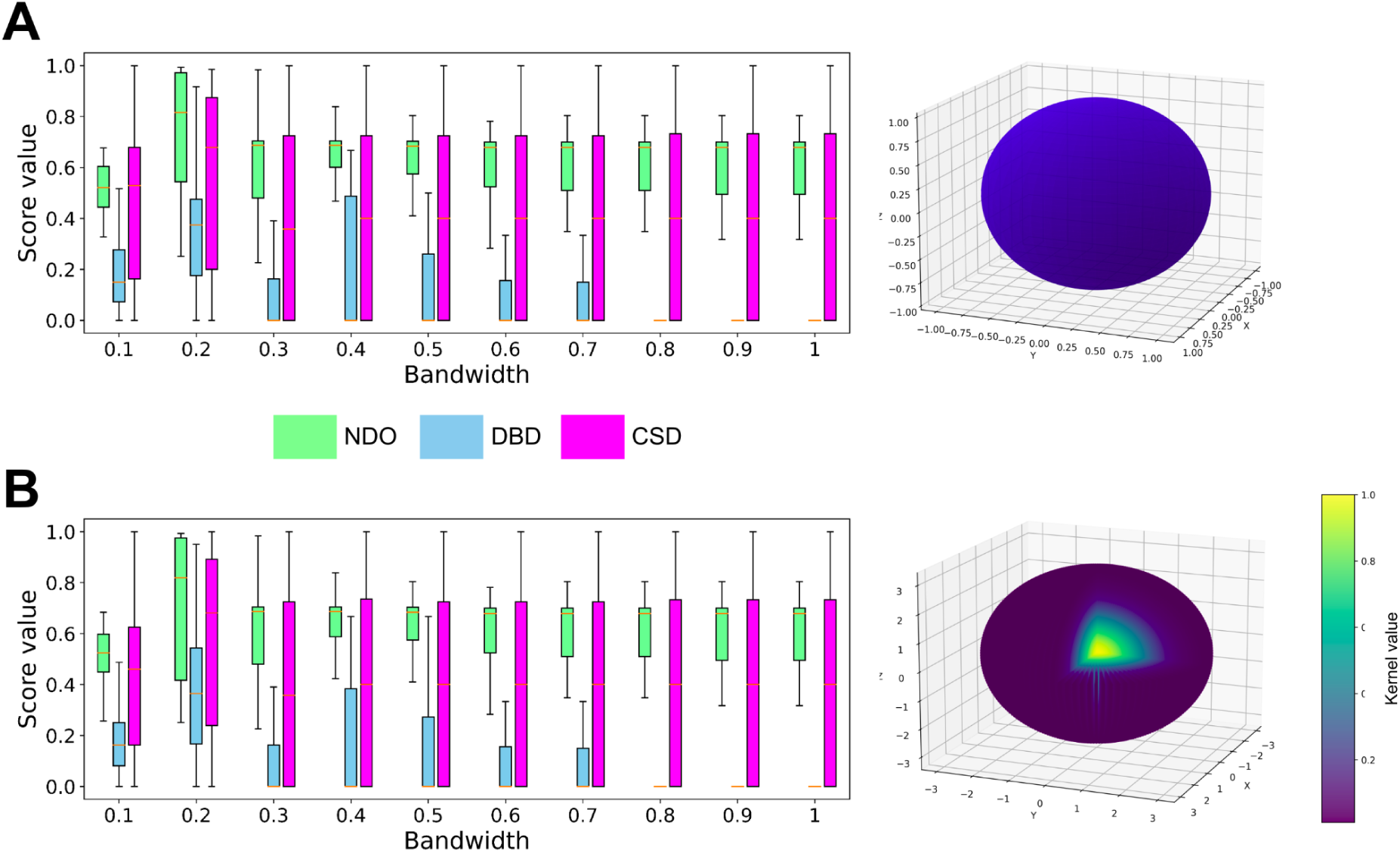
The NDO, DBD, and CSD values obtained with the Mean Shift clustering, using different hyperparameter values calculated on the subset of multi-domain RNA structures (*n*=50 entries) from the RNA3DHub set. For the bandwidth, 10 values were tested (expressed as a ratio of quantile), either using (**A**) a uniform kernel, or (**B**) a Gaussian kernel. For illustrative purposes, the corresponding kernel in the three-dimensional feature space is shown in the right panel; it is an ellipsoid in both cases. The kernel value is represented by a color, with a cutaway view for the Gaussian kernel.

### 3.3. Segmentation quality evaluation

We assessed the performance of RNA3DClust using three datasets of RNA 3D domains constructed for this study. For each dataset, a quantitative evaluation of the segmentation quality is provided, based on the five different metrics described in Section 2.4, including our newly developed CSD score (Fig. 5). This is followed by a qualitative evaluation, illustrating various examples of both correct and incorrect domain decompositions. The values presented correspond to the C3’-based segmentation (Tables S6 to S8), which yielded the highest overall performance; results for C1’ and C4’ are provided in Tables S9 to S11, and Tables S12 to S14, respectively.

**Figure 5.**
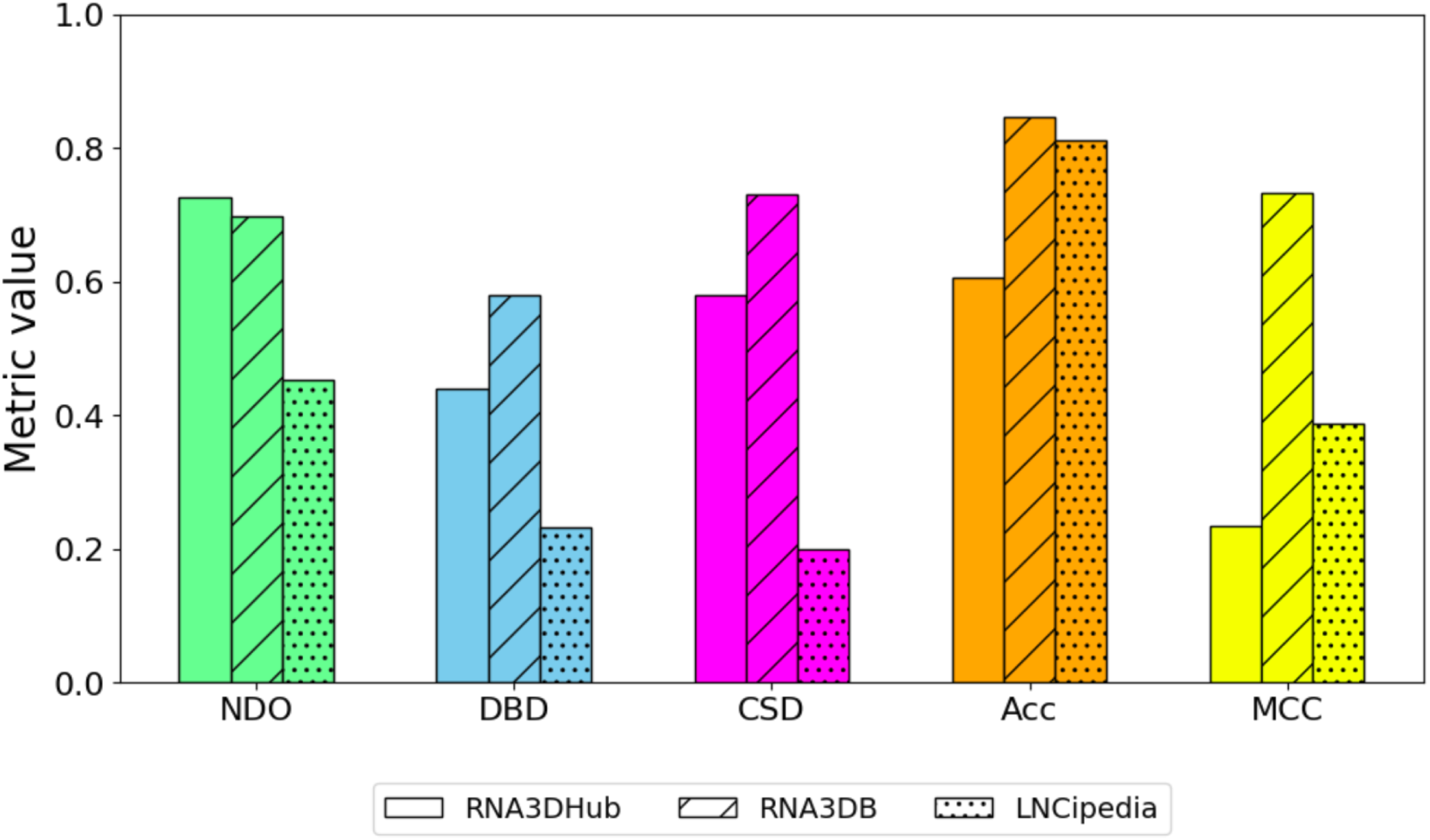
Values of the five performance metrics for the three RNA 3D structure datasets.

#### 3.3.1. Performance on the RNA3DHub and RNA3DB sets

For the 132 (single- and multi-domain) experimental RNA structures from the RNA3DHub set, the average NDO, DBD, and CSD values obtained were 0.726, 0.441, and 0.579, respectively (Fig. 5). For the 26 (single- and multi-domain) experimental RNA structures from the RNA3DB set, the average NDO, DBD, and CSD values obtained were 0.697, 0.580, and 0.730, respectively (Fig. 5). For indirect comparison, state-of-the-art protein domain parsers, like UniDoc [20] and FUpred [56], achieve average NDO scores of 0.812 and 0.804, respectively. Having a closer look at the results for these two datasets, we observed that RNA3DClust correctly classified single- and multi-domain structures in 80 of the 132 RNA3DHub cases (Table S6), and in 22 of the 26 RNA3DB cases (Table S7). This corresponds to an Acc of 60.61% and a MCC of 0.234 for RNA3DHub, and an Acc of 84.62% and a MCC of 0.732 for RNA3DB (Fig. 5). Again, these performance values may be considered in the light of those obtained by recent protein domain parsers, with UniDoc and FUpred achieving MCC of 0.804 and 0.799, respectively [20].

To understand the strengths and limitations of our method, we visualized the results produced by RNA3DClust for each of the RNA structures from the RNA3DHub and RNA3DB sets, and present in Fig. 6 eight cases: four correct domain assignments (Fig. 6A to 6D) and four incorrect ones (Fig. 6E to 6H). In the case of the *Escherichia coli* 16S rRNA (Fig. 6A), which has a two-domain structure, the post-clustering procedure behaved as expected, by removing all outlier regions and expanding the clusters, resulting in an almost perfect annotation. The same applies to the *Oryctolagus cuniculus* (European rabbit) 18S rRNA (Fig. 6B), where the empirically defined maximum linker length of 100 nt resulted in the removal of the outlier, thereby making the results consistent with the ground truth (with the exception of a relatively short and discontinuous additional domain). The synthetic single guide RNA (sgRNA) is a case of a two-domain structure with a “terminal” linker (Fig. 6C), for which RNA3DClust successfully found two clusters and one outlier, while a third spurious cluster was eliminated during post-processing. The case of the Taura syndrome virus internal ribosome entry site (IRES) RNA (Fig. 6D) also represents a two-domain structure, but without a linker, for which RNA3DClust computed a segmentation that perfectly matched the ground truth, thanks to the post-clustering procedure.

**Figure 6.**
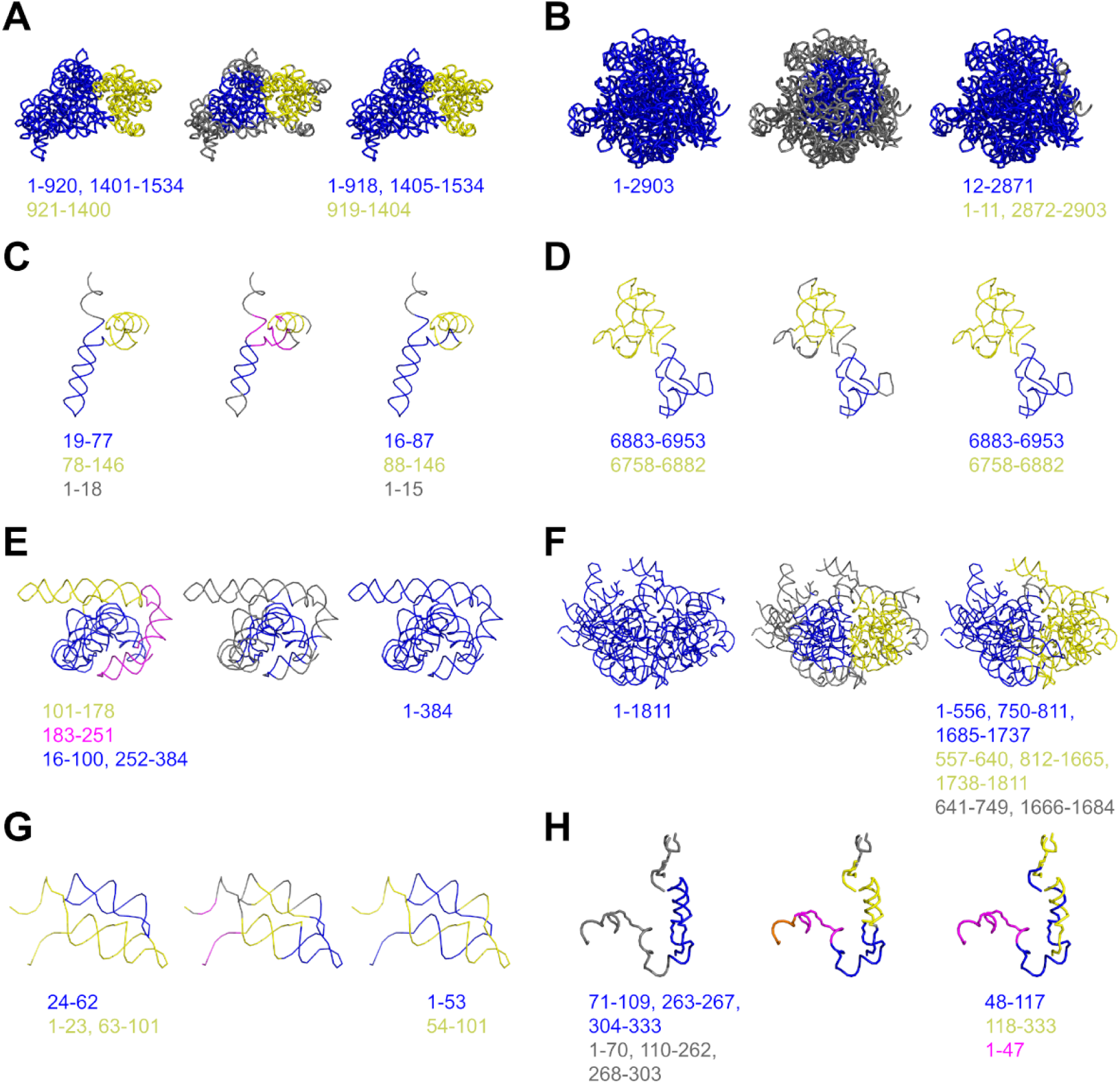
Examples of segmentations for experimental RNA 3D structures of the RNA3DHub and RNA3DB sets. For each RNA, the reference segmentation is presented on the left panel; on the middle and right panels, RNA3DClust results are presented, before and after the post-clustering procedure, respectively. Labeled clusters are colored in blue, yellow, and magenta, while outliers are in gray. Sequence positions of the domains are written below the structures (see also Supplementary Tables). (**A**) *E. coli* 16S rRNA (PDB entry: 4YBB, chain AA), (**B**) *O. cuniculus* 18S rRNA (7O7Y, chain A2), (**C**) Synthetic sgRNA (6JDV, chain B), (**D**) Taura syndrome virus IRES RNA (8EWC, chain EC), (**E**) *Tetrahymena* ribozyme (8TJX, chain N), (**F**) *T. aestivum* 40S rRNA (8R6F, chain A), (**G**) *E. siraeum* THF riboswitch (3SUX, chain X), (**H**) *S. cerevisiae* U3 snoRNA (6KE6, chain 3A).

The first incorrect case is the under-segmentation of the *Tetrahymena* ribozyme three-domain structure (Fig. 6E), which RNA3DClust erroneously annotated as a single domain. Mean Shift initially identified a cluster corresponding to part of the compact domain but classified the two helical regions as outliers. This case illustrates that the post-processing step can merge or remove clusters and outliers but cannot generate new ones, and is therefore limited to correcting under-fragmentation in Mean Shift’s output. However, the single-domain structure of the *Triticum aestivum* (common wheat) 40S rRNA (Fig. 6F) was over-segmented by Mean Shift into two clusters and an outlier, but the post-processing failed to merge or remove them. The *Eubacterium siraeum* THF riboswitch exemplifies a case in which a multi-domain RNA structure consists of distinct double-helical domains that are not well separated in space (Fig. 6G). As a result, Mean Shift groups portions of different domains into the same cluster, producing an incorrect delimitation that may not be corrected by post-processing. Finally, the U3 small nucleolar RNA (snoRNA) of *Saccharomyces cerevisiae*, which consists of a single discontinuous domain and a discontinuous linker region (Fig. 6H), was over-segmented by Mean Shift into four clusters and an outlier. Post-processing incorrectly removed the outlier and merged only two domains, yielding a decomposition into three continuous domains.

#### 3.3.2. Performance on the LNCipedia set

We then ran RNA3DClust on the lncRNA predicted structures from the LNCipedia set (*n*=69 entries). The average NDO, DBD, and CSD values obtained were 0.454, 0.232, and 0.200, respectively (Fig. 5). In the single- vs multi-domain classification (see Section 2.4.4), RNA3DClust correctly found the class in 56 of the 69 cases (Table S8), with a MCC of 0.388 (Fig. 5). These low values correspond to a decrease in performance, compared to those reported for the RNA3DHub and RNA3DB sets in the previous section. This may be attributed to the fact these two datasets were alternatively used for hyperparameter tuning, whereas the LNCipedia set served as a test set only. Additionally, the entries in the RNA3DHub and RNA3DB sets consist of native 3D structures, whereas those in the LNCipedia set are predictions, in which the hierarchical organization of RNA 3D structure may not be perfectly recreated. For completeness, the Acc value is presented in Fig. 5, but this metric is not well-suited to the LNCipedia set, due to the class imbalance (15 single-domain vs 54 multi-domain RNA structures).

Based on the segmentation score values (Table S8), we selected for illustration four cases of correct segmentation (Fig. 7A to 7D) and four cases showing the limitations of our method (Fig. 7E to 7H). In the four correct examples, we can see that the post-clustering procedure successfully extended clusters over outlier regions, making the output segmentation less fragmented. In the case of the regulatory and Alzheimer’s disease-related lncRNA EPB41L4A-AS1 (Fig. 7A), the post-processing step also removed a spurious cluster initially produced by Mean Shift, yielding the correct annotation of two domains and a discontinuous linker region. For the Arylacetamide Deacetylase Like 2 (AADACL2) antisense RNA 1 (Fig. 7B), the computed boundaries closely matched the ground truth. Although one of the delimited segments was incorrectly classified as a cluster rather than an outlier, this decomposition can still be considered correct. Interestingly, portions of two distinct double-helical domains were grouped into the same cluster by Mean Shift due to their spatial proximity, but the post-processing step subsequently separated them correctly. The apoptosis associated transcript in bladder cancer (AATBC) lncRNA has a single-domain structure (Fig. 7C), which was successfully identified by RNA3DClust, apart from a short 12-nt linker left by the post-clustering procedure. This again illustrates how the empirically defined maximum linker length of 100 nt enables the successful removal of outliers. A similar case was observed for the ACTN1 antisense RNA 1 (Fig. 7D), except that the spurious linker was entirely removed, resulting in a perfect single-domain assignment.

**Figure 7.**
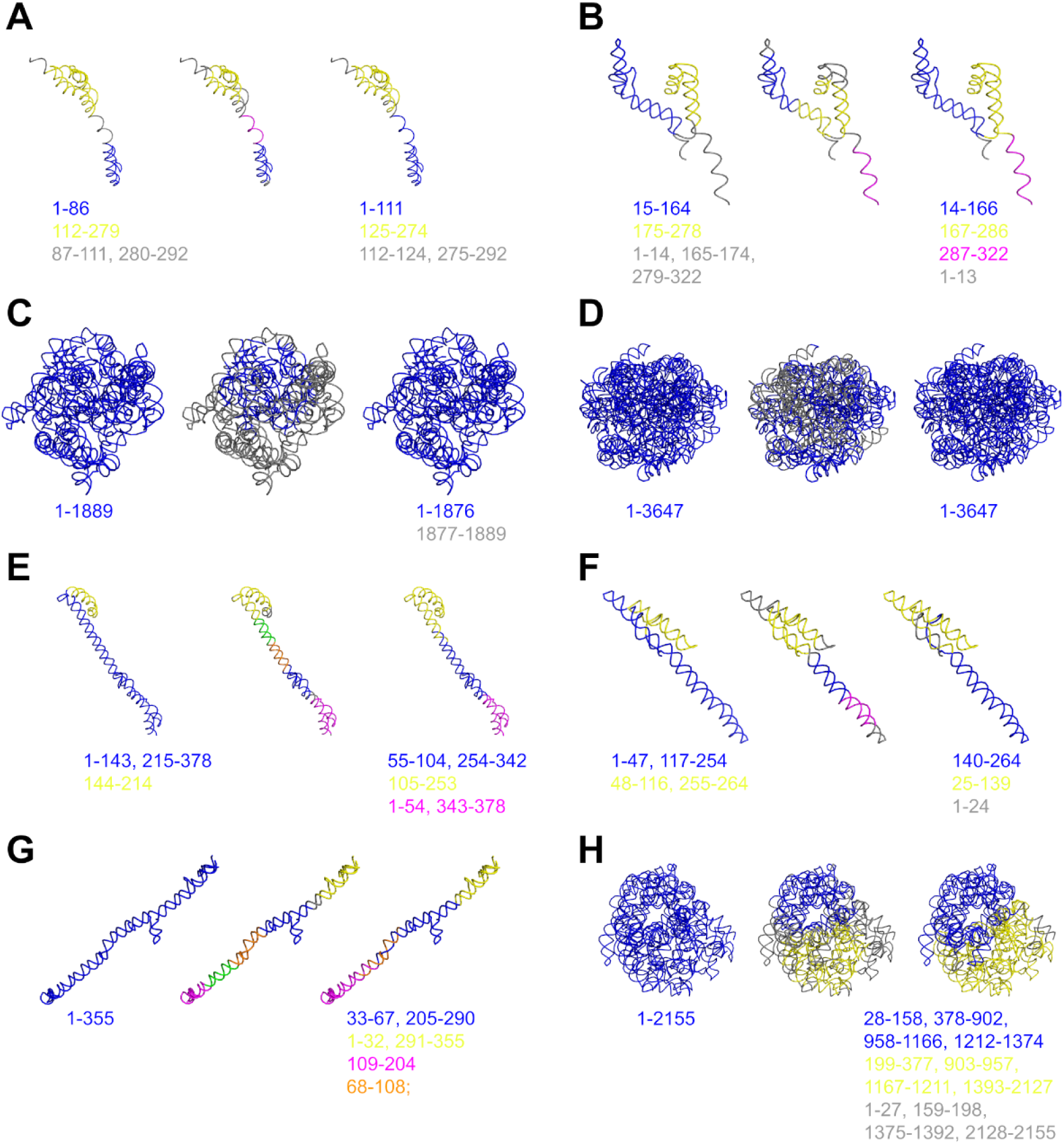
Examples of segmentations for predicted 3D structures of the LNCipedia set. For each lncRNA, the reference segmentation is presented on the left panel; on the middle and right panels, RNA3DClust results are presented, before and after the post-clustering procedure, respectively. Labeled clusters are colored in blue, yellow, orange and magenta, while outliers are in gray. Sequence positions of the domains are written below the structures (see also Table S8). The LNCipedia entries are: (**A**) EPB41L4A-AS1:6; (**B**) lnc-AADACL2-AS1:5; (**C**) AATBC:12; (**D**) ACTN1-AS1:14; (**E**) AADACL2-AS1:6; (**F**) FTX:15; (**G**) AATBC:2; (**H**) linc01136:5.

In proteins, 3D domain decomposition results in only two classes of substructures: domains and linkers. For this pioneering work on RNAs, we chose to include double helical segments into the domain class, rather than defining a third category. However, this particular type of domain poses a greater challenge for RNA3DClust than regular 3D domains, as their definition does not necessarily satisfy the spatial separation criterion. Moreover, when these straight double-helical substructures exceed a certain length, they tend to be over-fragmented by our method. This arises from the use of a fixed (rather than adaptive) bandwidth in Mean Shift, with the post-processing step subsequently failing to merge the artifactual clusters. Therefore, it is not surprising that the worst segmentation results were obtained on structures that are mainly composed of double helical segments. For example, another transcript of the AADACL2 antisense RNA 1 (Fig. 7E), which has a two-domain structure, was initially decomposed into five domains by Mean Shift, with parts of the two ground-truth domains incorrectly grouped within the same cluster. The post-processing step only partially corrected this over-segmentation by merging three clusters, while failing to separate the misgrouped domains. The next case, the FTX transcript XIST regulator (Fig. 7F), is similar, although the error made by RNA3DClust was less pronounced. In contrast, for the long intergenic non-protein coding RNA (lincRNA) 1136 (Fig. 7G), the severe over-fragmentation of its single-domain structure by Mean Shift was only minimally compensated for by post-processing, with just two of the five clusters being merged. Finally, another transcript of the AATBC lncRNA (Fig. 7H), despite having a compact structure, was over-segmented by Mean Shift into two domains and a linker, which the post-processing step failed to merge into a single-domain annotation.

### 3.4. Relevance to biological function and evolution

The present work introduces the idea of parsing spatially separate and compact regions from RNA 3D structures. In proteins, domains may also be defined based on their capacity to function or evolve in an independent manner. These functional and evolutionary criteria often produce the same domains than those delimited based on 3D geometry. However, the existence of such multiple criteria is sometimes a source of discrepancies among domain annotations [30]. Here, we wanted to address the question of the agreement between the clusters generated by RNA3DClust and the functional and evolutionary domains that are documented in the Rfam database. Among the 128 Rfam consensus annotations defined in Section 2.5.3 (122 from RNA3DHub and 6 from RNA3DB), we present eight representative examples: four cases of correct segmentation by RNA3DClust (Figs. 8A to 8D) and four cases of failure (Figs. 8E to 8H). In each category, we selected two structures from the RNA3DHub set (Figs. 8A, 8B, 8E, and 8F) and two structures from the RNA3DB set (Figs. 8C, 8D, 8G, and 8H). Details of the 128 cases are available in Tables S4 and S5. Note that these tables identify domains using the residue numbers from the PDB file, while Fig. 8 shows their positions along the RNA sequence. Because PDB numbering may differ from sequence numbering (e.g., if the PDB entry starts at a residue other than 1, or if some residues are missing from the structure), the two representations may not align perfectly.

**Figure 8.**
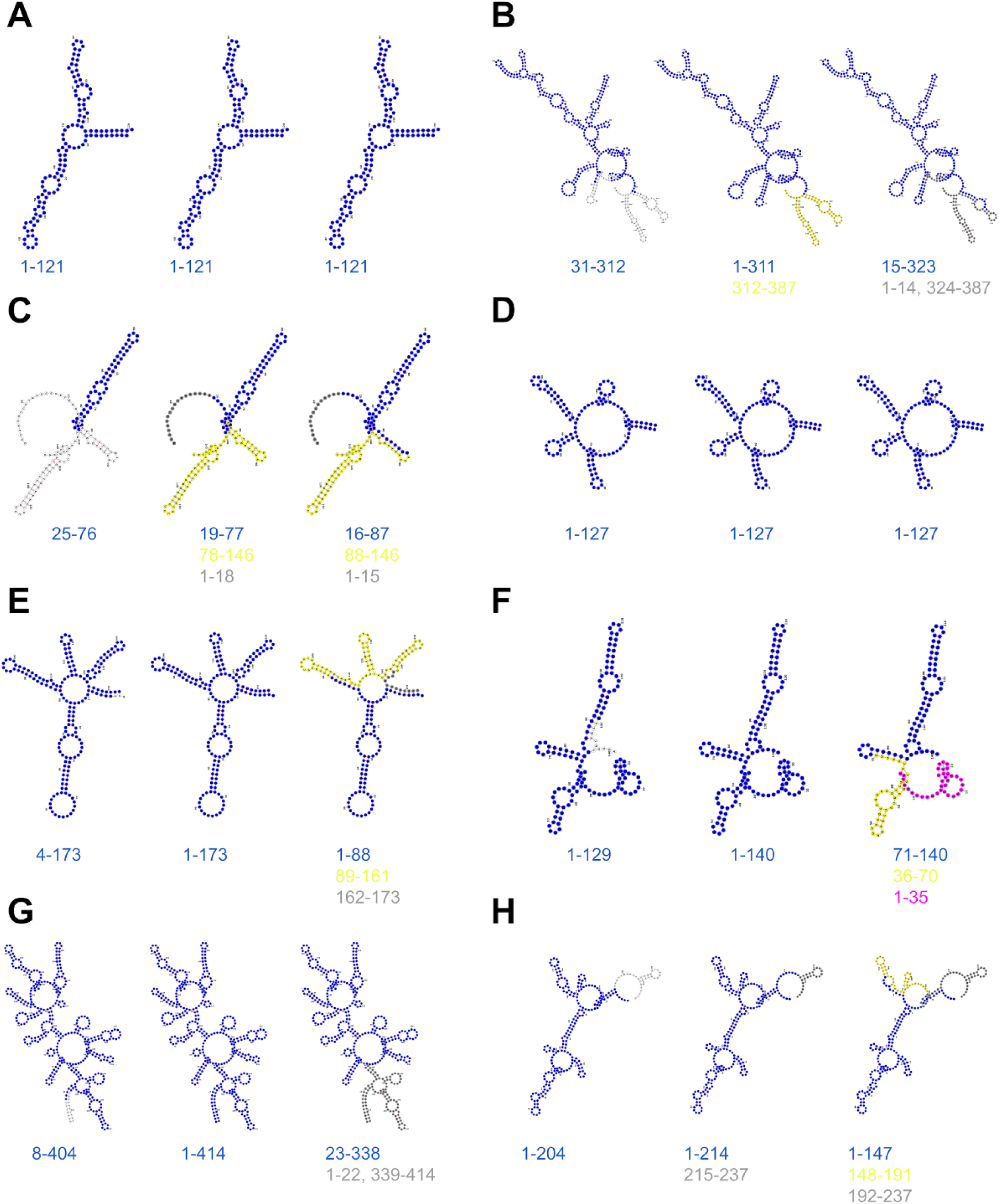
Examples of RNA domain delimitations showing consensus between the Rfam database (left subpanels) and visual inspection of 3D geometry (middle subpanels), with the decompositions produced by RNA3DClust displayed in the right subpanels. The PDB and Rfam entries are: (**A**) *S. cerevisiae* 5S rRNA with its 5S rRNA family domain (PDB: 8CCS, chain BB; Rfam: RF00001); (**B**) *T. thermophila* ribozyme with its group I catalytic intron domain (9CBX, chain N; RF00028); (**C**) Synthetic sgRNA with its CRISPR RNA direct repeat element domain (6JDV, chain B; RF01344); (**D**) *R. gauvreauii* yjdF riboswitch with its azaaromatic riboswitch aptamer domain (8UIW, chain R; RF01764); (**E**) *T. maritima* lysine riboswitch with its lysine riboswitch aptamer domain (3DIL, chain A; RF00168); (**F**) *H. sapiens* GlmS ribozyme with its glmS glucosamine-6-phosphate activated ribozyme aptamer domain (2NZ4, chain Q; RF00234); (**G**) *B. stearothermophilus* ribonuclease P RNA with its bacterial RNase P class B domain (2A64, chain A; RF00011); (**H**) HCV IRES RNA with its hepatitis C virus IRES domain (7SYS, chain z; RF00061).

Out of the 122 domain annotations in our Rfam consensus set (Section 2.5.3), RNA3DClust correctly delimited 88 (IoU ≥ 0.8) and incorrectly delimited 34, corresponding to an accuracy of 72.1% and an average IoU of 0.837 ± 0.207. The 5S rRNA of *S. cerevisiae* is annotated in Rfam as belonging to the RF00001 family over its entire length (Fig. 8A), which agrees with our assignment of a single 3D domain based on structural compactness. In this case, RNA3DClust successfully grouped all residues into one cluster. For the *T. thermophila* ribozyme, Infernal identified a group I catalytic intron structure (RF00028) that spans most of the 387-nt chain (Fig. 8B). Visual inspection revealed a clear 3D-structural distinction between most of the annotated Rfam domain and the remainder of the structure, resulting in the assignment of two 3D domains. Here, RNA3DClust correctly delineated the group I catalytic intron domain (but misclassified the smaller 3D domain as a discontinuous linker region).

Out of the six domain annotations showing consensus between the RNA3DB set and Rfam, only two were correctly delimited by RNA3DClust. The first is the synthetic sgRNA, which contains a CRISPR RNA direct repeat element with a stem-loop (Fig. 8C). By visual inspection, this substructure is clearly distinct from the rest of the macromolecule, leading us to assign two 3D domains. RNA3DClust successfully captured this distinction, assigning the stem-loop to a separate cluster, with boundary differences of only a few residues. The second case is the *Ruminococcus gauvreauii* yjdF riboswitch (Fig. 8D), which, although different from the 5S rRNA described above, is likewise annotated as a single Rfam domain (the azaaromatic riboswitch aptamer domain; RF01764) and visually forms a compact 3D domain. Here too, RNA3DClust correctly grouped all residues into one cluster. Overall, such examples of agreement between the segmentations based on 3D geometry and those based on function and evolution show the potential of RNA3DClust to delimit biologically relevant RNA substructures. More importantly, it highlights the pertinence of decomposing RNA into 3D structural domains.

Besides the successful cases described above, RNA3DClust also failed to recover certain Rfam domains. For the *Thermotoga maritima* lysine riboswitch, the entire structure corresponds to an Rfam domain (RF00168) (Fig. 8E), consistent with our compactness-based assignment of a single 3D domain. However, RNA3DClust over-segmented it into two clusters, the larger one having a low IoU of 0.5 with the Rfam domain. Similar over-segmentation errors explain the next three failures: two spurious clusters in the *Homo sapiens* GlmS ribozyme (Fig. 8F), a spurious outlier region in the *Bacillus stearothermophilus* ribonuclease P RNA (Fig. 8G), and both a spurious cluster and an outlier region in the hepatitis C virus IRES RNA (Fig. 8H). All these cases resulted in IoU values < 0.8 (in Fig. 8G, the reported IoU of 0.8 is due to rounding in Table S5). Importantly, although RNA3DClust failed to detect both the Rfam and 3D domains in these examples, the resulting clusters nevertheless corresponded to substructures with distinct secondary structure features. This suggests that RNA3DClust’s segmentations may still be meaningful at finer levels of hierarchical organization, such as RNA motifs.

### 3.5. Comparison of the scoring functions

#### 3.5.1. Score variations across datasets

In this work, we introduced a new segmentation quality score, the CSD, and used the two most cited ones, the NDO and the DBD. For these metrics, we observed values ranging from 0.0 to 1.0, except two negative NDO values (−0.225 for 6KE6 chain 3A and −0.029 for lnc-NKX2-3-1:2, in the RNA3DB and LNCipedia sets, respectively). As shown in Fig. 9 for the multi-domain structures of the three datasets, the NDO values tend to be relatively high, while the DBD values are typically lower, with average scores of 0.592 for NDO and 0.311 for DBD. This is due to the DBD being stricter in penalizing margin deviations when measuring boundary distances: even if the computed and true domains share a high percentage of overlapping nucleotides (high NDO), their boundaries can still be distant, resulting in a low DBD. Interestingly, the CSD, which we designed as a compromise between the NDO and the DBD, was in between these two metrics on the RNA3DHub set, and was close to either the NDO on the RNA3DB set, or the DBD on the LNCipedia set (see also Fig. 5). Because this behavior is similar for both single- and multi-domain RNAs, here we restrict our analysis to multi-domain structures; results for single-domain RNAs are provided in Fig. S4.

**Figure 9.**
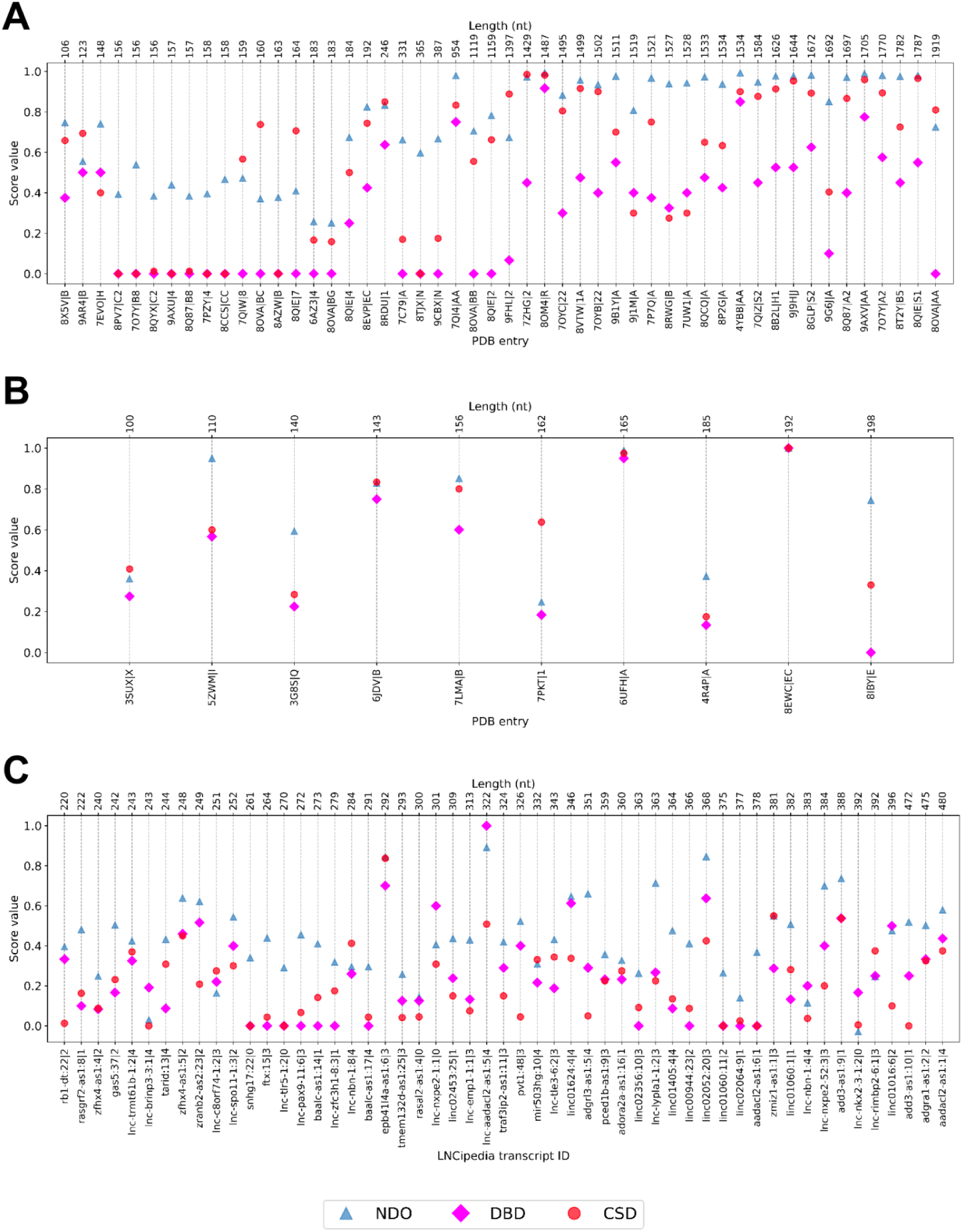
The NDO, DBD, and CSD values obtained for the multi-domain RNA 3D structures from (**A**) RNA3DHub, (**B**) RNA3DB and (**C**) LNCipedia dataset. The samples are sorted by ascending RNA length, from left to right of the figure. Detailed score values are provided in Tables S6 to S8.

The three scores vary similarly across the three datasets, with all metrics consistently indicating higher performance on the RNA3DB set and lower performance on the LNCipedia set (Fig. 5). The sole exception is the NDO, for which the difference between RNA3DHub and RNA3DB is only marginal. This apparent ranking of dataset difficulty can be explained by differences in their respective contents, as well as by the design of the scoring functions. Although RNA3DHub and RNA3DB have the same class balance (∼62% single- and ∼38% multi-domain), they differ markedly in RNA length, with entries in RNA3DHub being much longer (Figs. 9A and 9B). On these large RNA structures, the risk of over-segmentation is higher, which is penalized by all metrics except NDO. Even when the number of 3D domains is correct, the boundary distance tends to be greater in larger structures, leading to stronger penalties from DBD and, to a lesser extent, CSD. However, the lengths of LNCipedia entries are comparable to those in RNA3DB (Fig. 9C), yet LNCipedia appears to be the most challenging dataset. This may be due to LNCipedia containing more double-helical 3D domains (e.g., Figs. 7E to 7G), which RNA3DClust struggles to segment correctly—as an additional factor to the “predicted-versus-experimental structure” one mentioned in Section 3.3.2. The same qualitative explanation also accounts for RNA3DHub being more difficult than RNA3DB, as RNA3DHub contains a larger proportion of ambiguous double-helical cases (e.g., Fig. 6E).

#### 3.5.2. Correlation analysis

Although our new CSD score was inspired by the NDO and DBD, the three scoring functions rely on different approaches to assess the segmentation quality of RNA 3D structures (Section 2.4). Here, we investigated the difference in the information they actually capture, by studying the pairwise relationships between their values. Thus, for each pair of scores, we generated a scatter plot from the results obtained on the RNA3DHub (Figs. 10A to 10C), RNA3DB (Figs. 10D to 10F), and LNCipedia (Figs. 10G to 10I) sets. Strikingly, across the three datasets, most of the outlier points correspond to single-domain structures (red dots in Fig. 10). This arises from the fact that the DBD can only take the values 0 or 1 for single-domain RNAs. As expected, the CSD shows a hybrid behavior: it often tends toward these extreme values but can also take intermediate values, as can the NDO. To properly estimate correlations, we therefore restricted the calculations to the multi-domain structures of each dataset.

**Figure 10.**
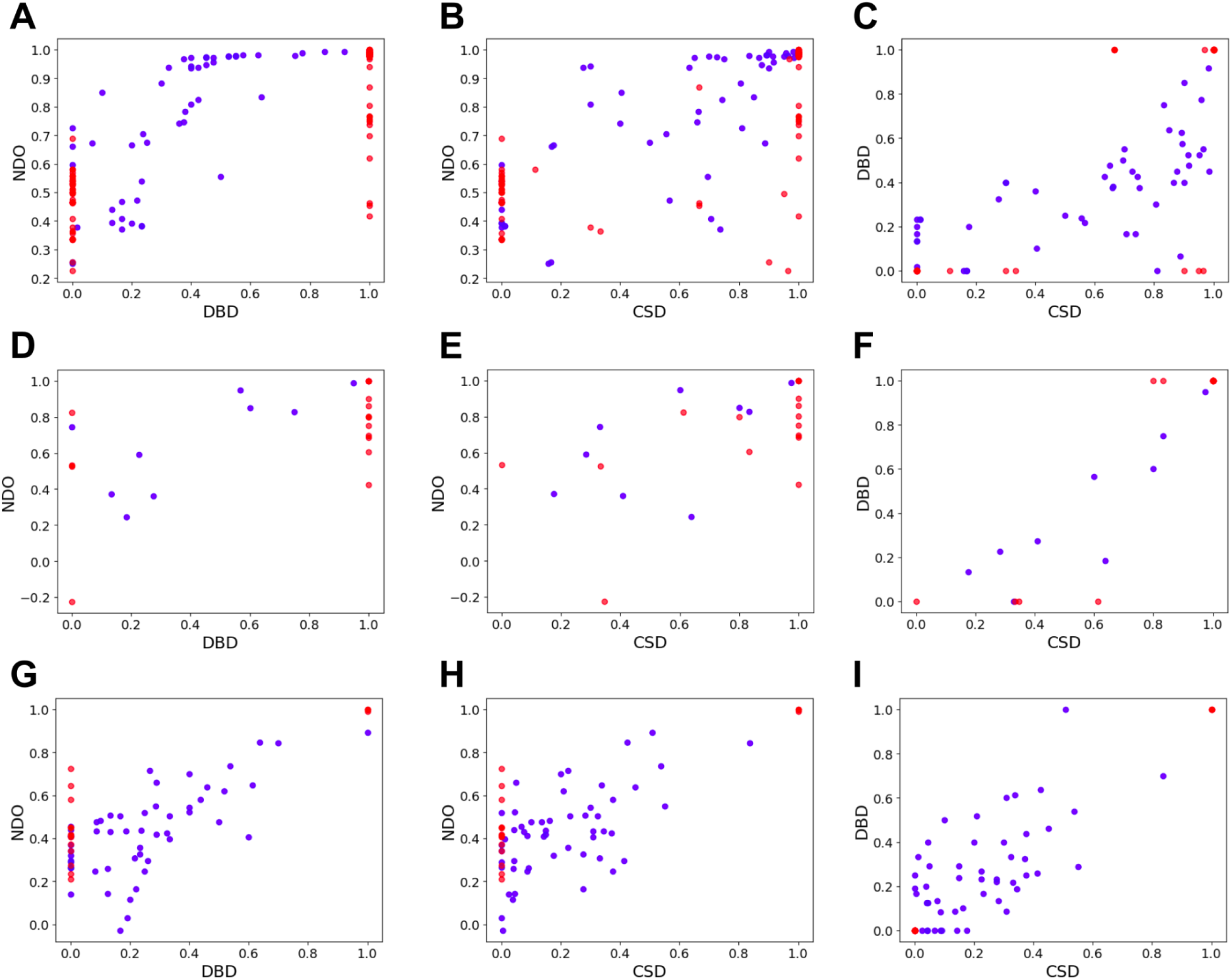
Scatter plots showing the relationship between each pair of segmentation quality scores, for the RNA 3D structures from (**A**, **B**, **C**) RNA3DHub (*n*=132), (**D**, **E**, **F**) RNA3DB (*n*=26) and (**G, H, I**) LNCipedia (*n*=69). Entries annotated as single- and multi-domain structures are represented by red and blue dots, respectively.

For all datasets, all pairs of scoring functions exhibit strong positive linear correlations, with the weakest *ρ* value being 0.590 for the CSD-NDO pair on the multi-domain subset of LNCipedia (*n*=54 entries; blue dots in Fig. 10H), and the strongest *ρ* value being 0.909 for the CSD-DBD pair on the RNA3DB subset (*n*=10 entries; blue dots in Fig. 10F). In terms of statistical significance, all associated *p*-values were below 10^−2^, except for the CSD-NDO pair on the RNA3DB subset (*p*=3.67 × 10^−2^; *ρ*=0.663; Fig. 10E), likely due to the small sample size. The ranking of score pairs by correlation strength differs across datasets: the DBD-NDO pair is the most correlated only for RNA3DHub and LNCipedia, whereas the CSD-NDO pair is the least correlated only for RNA3DB and LNCipedia. This dataset-dependent behavior is consistent with the observations reported in previous sections, which we attributed to differences in dataset content. Overall, these correlation values demonstrate that each score is designed to capture different—yet congruent—aspects of RNA segmentation quality, thus providing complementary insights. Finally, it is worth noting that the DBD-NDO pair for RNA3DHub appears to display a non-linear but monotonic relationship (Fig. 10A). Since fitting a more appropriate model would only increase the correlation strength, this observation does not alter the above conclusions.

## 4. CONCLUSION AND PERSPECTIVES

This work introduced RNA3DClust, a computational tool for delimiting compact and spatially separate domains in RNA 3D structures. The method operates by clustering C3’ atoms using the Mean Shift algorithm, followed by a dedicated post-clustering procedure. Despite the limited amount of available data, RNA3DClust demonstrated promising performance at decomposing RNA 3D structures, notably in a biologically meaningful way. Nevertheless, as a pioneering work, several avenues of improvement can be identified. One potential enhancement is the implementation of an adaptive bandwidth, *i.e.* a flexible window size that varies proportionally to the distribution of points at each iteration of the Mean Shift algorithm. Of note, this would define (i) the bandwidth as a model parameter, compared to the current version where the bandwidth is a hyperparameter, as it is not learned directly from the data but, instead, fixed prior to running the algorithm, and (ii) the learning pipeline as unsupervised, as it would no longer involve the initial stage of tuning the Mean Shift kernel (bandwidth and type) using labeled data. The choice of Mean Shift was made from among the clustering algorithms most commonly-used and readily-available in scikit-learn—which has recently proven adequate for protein 3D domain decomposition [57]. However, alternative approaches could also be considered, such as graph-based methods, which have already shown effectiveness in decomposing protein 3D structures [58]. Finally, future work could explore leveraging the domain segmentation produced by RNA3DClust to improve RNA 3D structure prediction by training models on datasets of 3D domains, or to assess the quality of 3D models of long RNAs, akin to methods previously developed for protein structures [59].

## ACKNOWLEDGEMENTS

We thank the anonymous reviewers for their helpful comments. The authors gratefully acknowledge the financial support of the Ambassade de France au Vietnam (Bourses France Excellence, bc0180).

## Notes

### Competing Interest Statement

The authors have declared no competing interest.

### Summary of Updates

The datasets used to train and benchmark the method have been changed. Consequently, the Results and Discussion section has also been modified. Multiple paragraphs of explanations have been added to the Methods section. The title has been changed as the method actually qualifies more as "semi-supervised" than "unsupervised".

