## Supplementary figures for "Semi-supervised segmentation of RNA 3D structures using density-based clustering"

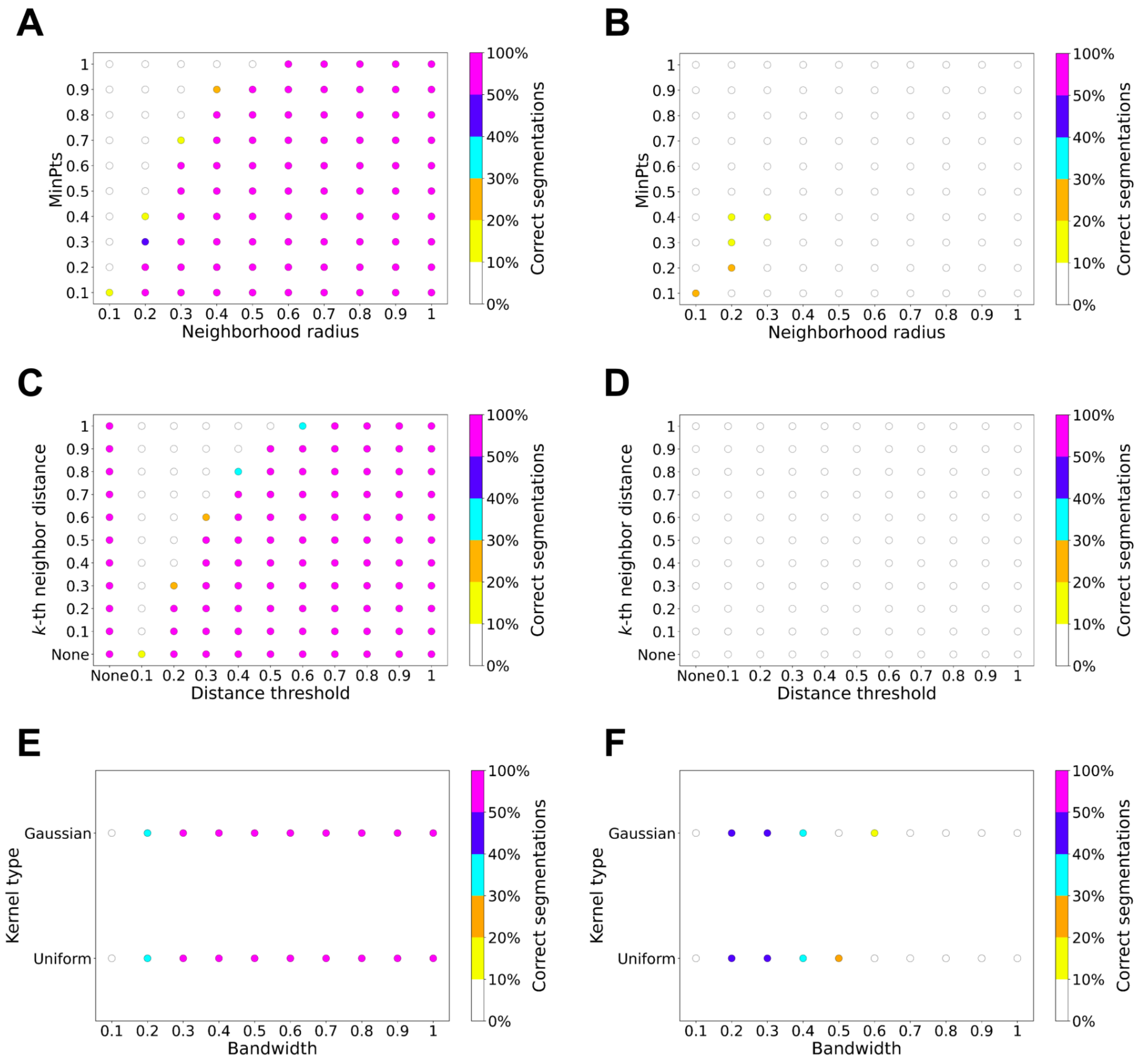


**Figure S1.** Scatter plots representing the search for hyperparameter values that return a correct segmentation in terms of number of domains, for (**A**, **B**) DBSCAN, (**C**, **D**) HDBSCAN and the hybrid approach, and (**E**, **F**) Mean Shift. The computations were performed on the 26 experimental RNAs structures from the RNA3DB set: (**Left panels**) The subset of 16 single-domain annotations; (**Right panels**) The subset of 10 multi-domain annotations. The color of each dot represents the percentage of cases with the right number of clusters, as shown by the color scale.

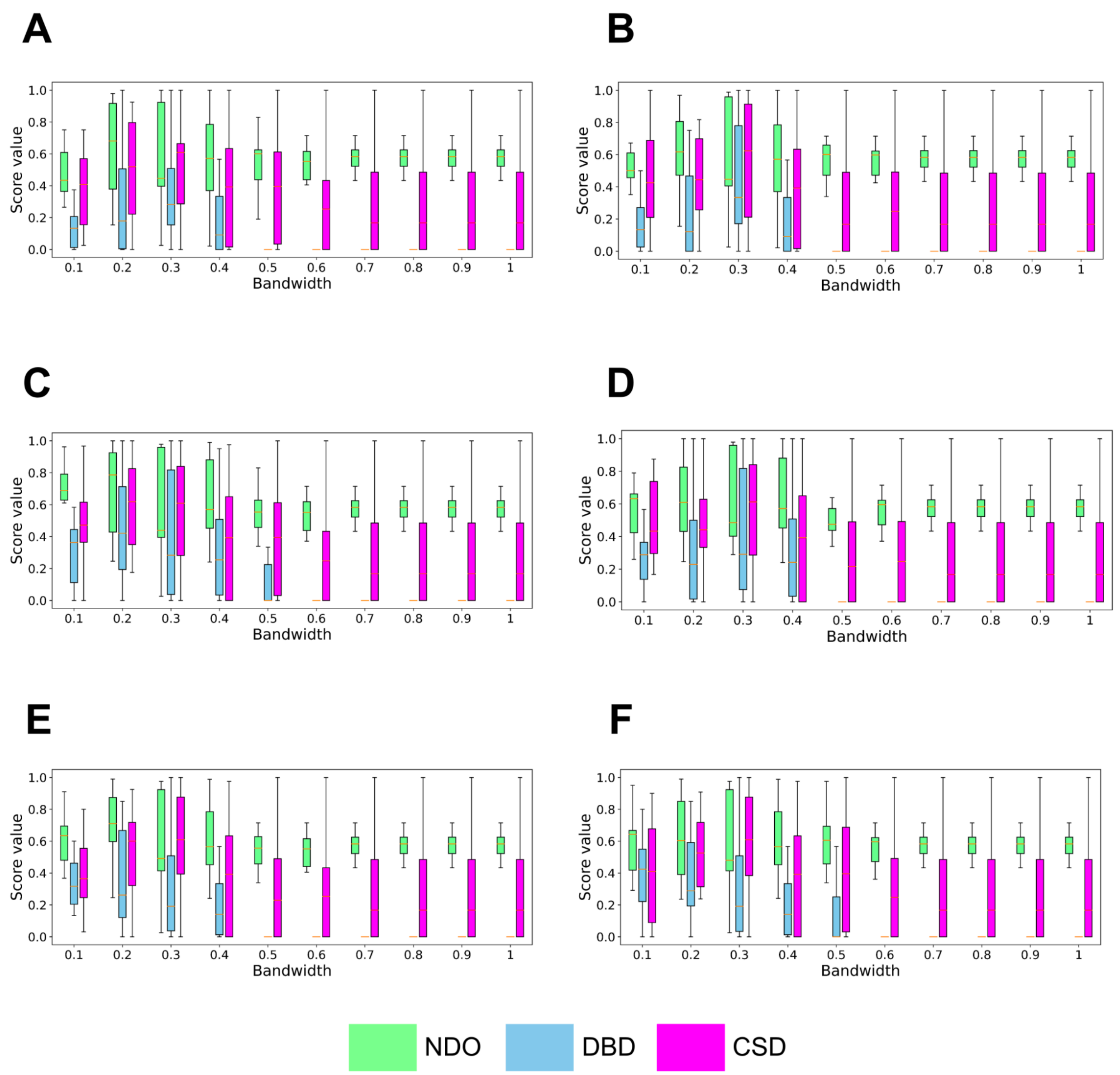


**Figure S2.** The NDO, DBD, and CSD values obtained with the Mean Shift clustering, using different hyperparameter values calculated on the subset of multi-domain RNA structures (*n*=10 entries) from the RNA3DB set. For the bandwidth, 10 values were tested (expressed as a ratio of quantile). Atomic representation and kernel type: (**A**) C1'-only + uniform kernel; (**B**) C1'-only + Gaussian kernel; (**C**) C3'-only + uniform kernel; (**D**) C3'-only + Gaussian kernel; (**E**) C4'-only + uniform kernel; (**F**) C4'-only + Gaussian kernel.

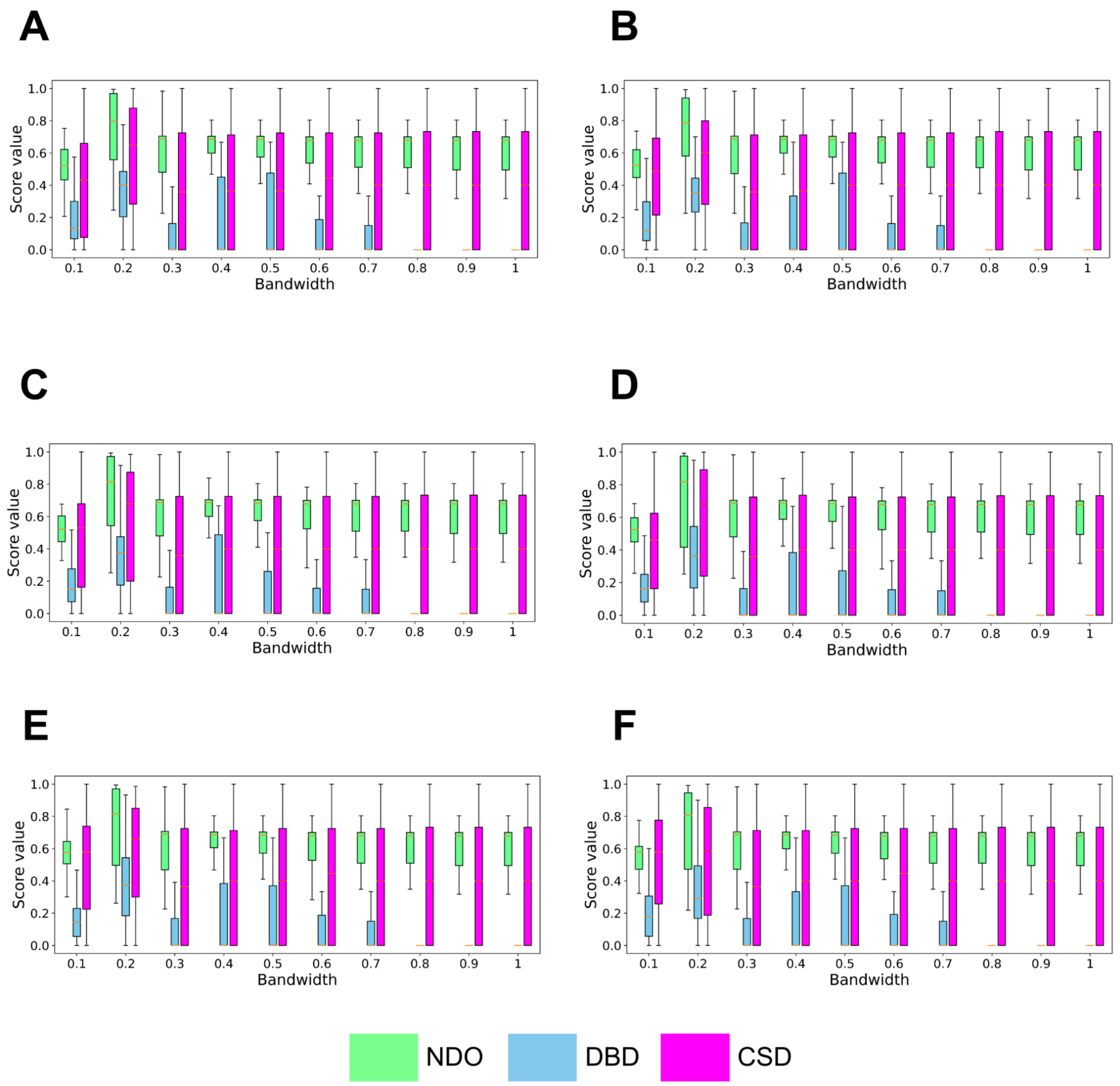


**Figure S3.** The NDO, DBD, and CSD values obtained with the Mean Shift clustering, using different hyperparameter values calculated on the subset of multi-domain RNA structures (*n*=50 entries) from the RNA3DHub set. For the bandwidth, 10 values were tested (expressed as a ratio of quantile). Atomic representation and kernel type: (**A**) C1'-only + uniform kernel; (**B**) C1'-only + Gaussian kernel; (**C**) C3'-only + uniform kernel; (**D**) C3'-only + Gaussian kernel; (**E**) C4'-only + uniform kernel; (**F**) C4'-only + Gaussian kernel.

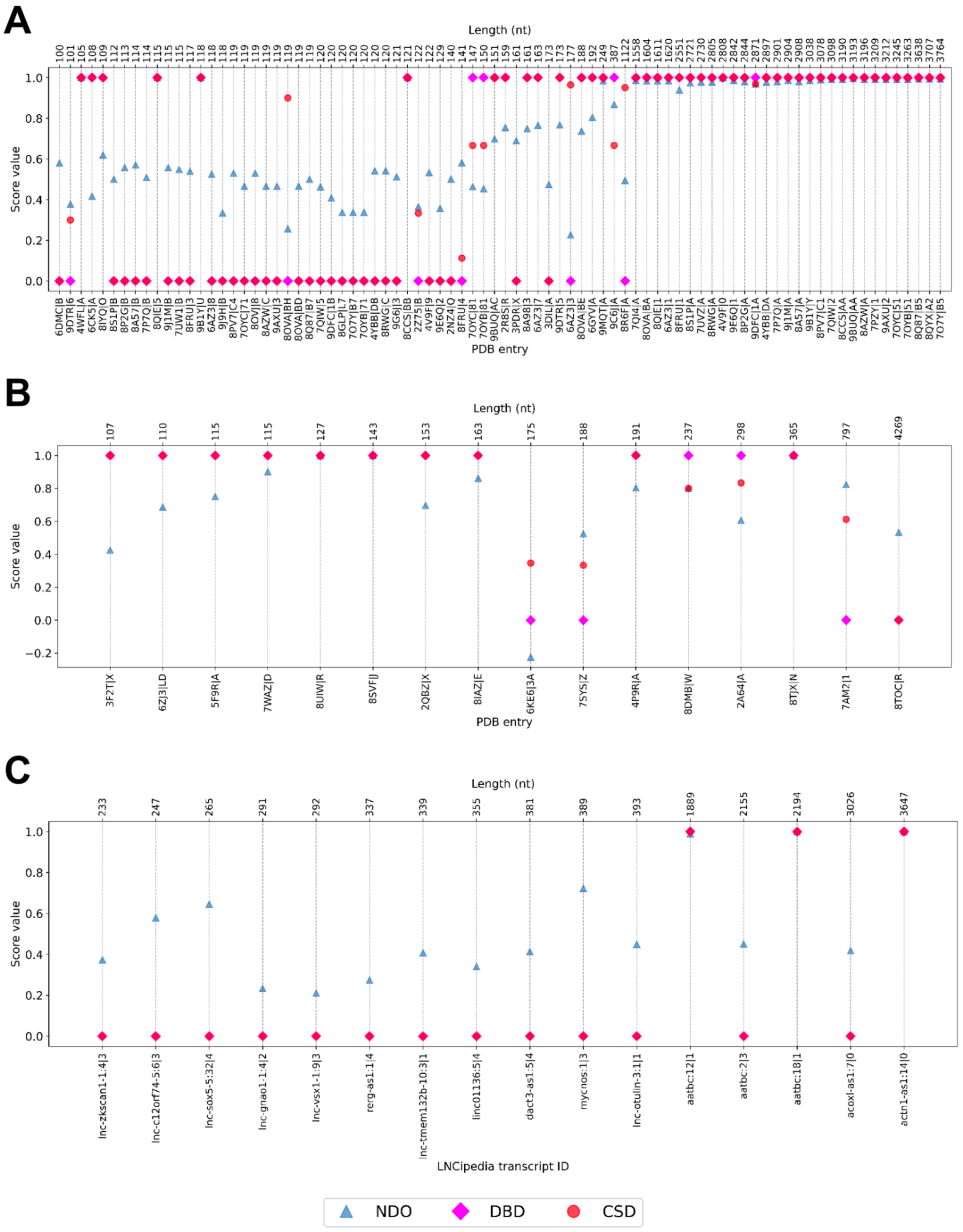


**Figure S4.** The NDO, DBD, and CSD values obtained for the single-domain RNA 3D structures from (**A**) RNA3DHub, (**B**) RNA3DB and (**C**) LNCipedia dataset. The samples are sorted by ascending RNA length, from left to right of the figure. Detailed score values are provided in Tables S6 to S8.
